# Atypical GPCR Activation Resolved by Nanobody Engineering

**DOI:** 10.64898/2026.01.10.698836

**Authors:** Roman R. Schlimgen, Shawn E. Jenjak, Alexa De La Sancha, Joy Darcis, Christian B. Billesbølle, Linda J. Olson, Francis C. Peterson, Martine J. Smit, Aashish Manglik, Martyna Szpakowska, Andy Chevigne, Brian F. Volkman

## Abstract

G protein-coupled receptors (GPCRs) are the largest family of clinically targeted proteins, yet most therapeutics target a narrow subset of structurally well-behaved receptors. The atypical chemokine receptor ACKR3 defies canonical models, displaying broad ligand recognition, high basal activity, and resistance to inhibition. Using engineered nanobodies, cryo-EM, NMR, and structure-guided pharmacology, we uncover an unconventional activation mechanism in ACKR3 that challenges established paradigms of GPCR activation. We find that receptor activity is controlled by changes in extracellular pocket volume rather than conformational rearrangements in conserved microswitches, and an expanded aromatic cluster at the intracellular transducer binding pocket stabilizes the active state. These findings redefine how GPCRs can be modulated and open new strategies for targeting pharmacologically intractable receptors.

## Introduction

G protein–coupled receptors (GPCRs) are among the most successful drug targets in medicine, with over one-third of FDA-approved therapeutics acting on members of this superfamily^1^. However, this frequently cited statistic obscures a critical reality: the vast majority of these drugs target a narrow set of structurally tractable and pharmacologically well-behaved receptors. In fact, over 70% of GPCRs are therapeutically unexploited and remain poorly characterized^1,2^. This disparity suggests that the prevailing mechanistic framework of GPCR activation, built largely around a limited set of prototypical receptors, may fail to account for broader structural and functional diversity across the superfamily^3^.

The current paradigm, in which a unique ligand binds the orthosteric pocket and engages conserved GPCR microswitches ^4,5^, fails to explain how chemically diverse ligands can bind a single receptor and trigger equivalent signaling outcomes. A striking example is the atypical chemokine receptor ACKR3. Initially identified as a scavenger for two chemotactic cytokines, ACKR3 is now known to bind at least 13 endogenous ligands across four distinct families (Supplementary Table 1)^6–11^. These ligands span a wide range of molecular weights and scaffolds (Figure 1a), yet all engage the receptor with high affinity and induce β-arrestin recruitment but not G protein activation. Promiscuous GPCRs like ACKR3 expose the limits of a one-size-fits-all model and highlight the need for alternative mechanisms that govern receptor function outside classical paradigms.

**Figure 1.**
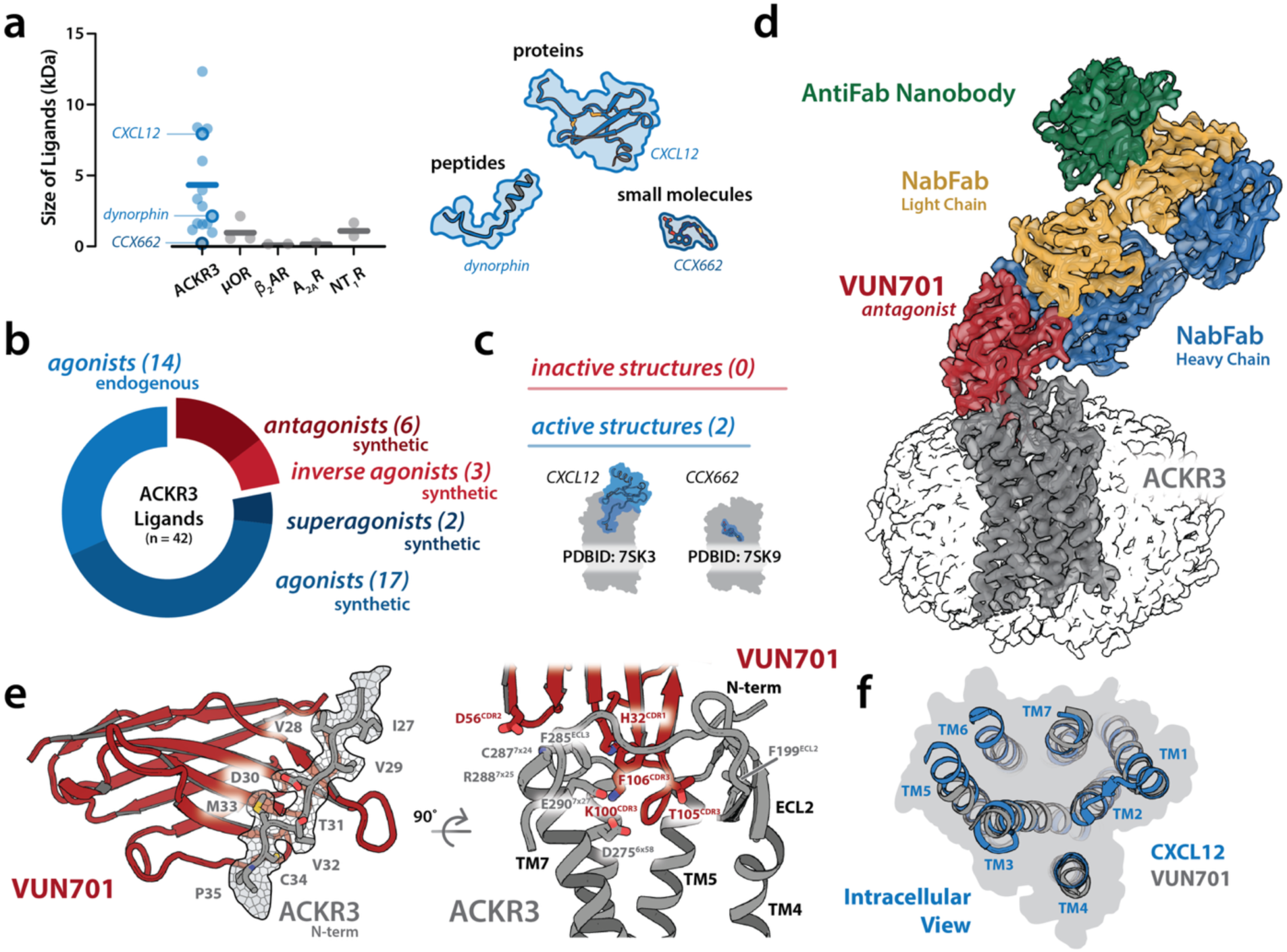
Structural basis of ACKR3 ligand recognition and nanobody engagement. a) Molecular weight distribution of endogenous ligands for ACKR3 and other prototypical GPCRs, illustrating ACKR3’s ability to accommodate ligands of diverse sizes (Supplementary Table 1). The synthetic ACKR3 agonist CCX662 is also plotted to illustrate the size of this scaffold. Right: Cartoon depiction of the scaffold diversity among ACKR3 ligands, spanning proteins (e.g., CXCL12), peptides (e.g., Dynorphin), and small molecules (e.g., CCX662). b) Summary of the complete pharmacological landscape of ACKR3, including 32 activating ligands (17 synthetic, 13 endogenous agonists, 2 synthetic superagonists) and only 9 inactivating ligands (6 synthetic antagonist and 3 synthetic inverse agonist) (Supplementary Table 2). c) The structures of CXCL12 and CCX662 agonist-bound complexes illustrate the receptor’s structural tolerance for divergent binding poses, but lack of structural information for the inactive state. d) Cryo-EM structure of ACKR3 (gray) in complex with the antagonist nanobody VUN701 (red). To facilitate cryo-EM particle alignment, the complex includes a nanobody-binding Fab (NabFab: light chain, gold; heavy chain, blue) and an anti-Fab nanobody (green). The receptor was imaged in an LMNG micelle (transparent surface). e) Left: VUN701 engages the N-terminal region of ACKR3, forming defined interactions with residues C26–P35 (gray – cryo-EM density shown as mesh surface), enhancing binding specificity and affinity. Right: VUN701 forms orthosteric interactions with ACKR3 through its CDR3 residues K100, T105, and F106. Additional contacts with ECL3 are made with H32 (CDR1) and D56 (CDR2). f) Intracellular view of the CXCL12-bound and VUN701-bound ACKR3 structures highlights the differences in intracellular resolution. While the CXCL12-bound structure shows a well-defined intracellular cavity with resolved TM5, TM6 and TM7 positions, the VUN701-bound structure lacks continuous density in these regions, precluding modeling of the intracellular face and resolving conformational changes.

ACKR3’s broad ligand recognition, together with drug discovery efforts that repeatedly yielded activating rather than inhibitory compounds, contributed to the prevailing view that ACKR3 is intrinsically activation-prone and resistant to pharmacological inhibition (Figure 1b, Supplementary Table 2)^12^. Indeed, several early small molecules, though potent competitors of CXCL12, were later revealed to be agonists, and several antagonists of the related receptor CXCR4 also acted as ACKR3 agonists^13–16^. The dearth of effective antagonists or inverse agonists has long hindered mechanistic and structural insight into ACKR3 activation (Figure 1c). However, recent efforts have yielded the first bona fide ACKR3 antagonists and inverse agonists, providing long-awaited tools to probe the receptor’s full activation landscape^17–20^.

Using one such antagonist, a 12 kDa nanobody (VHH) called VUN701, we determined the structure of ACKR3 and observed unexpected conformational heterogeneity dominated by an active-like state. Notably, the orthosteric pocket in this nominally inactive structure was more compact than in the structure of ACKR3 bound to its chemokine agonist CXCL12, in contrast to the pocket expansion typically observed for inactive-state GPCRs. To further explore this, we altered VUN701 to create agonists and inverse agonists. Structural comparisons of the ACKR3-nanobody complexes revealed an activation mechanism independent of canonical GPCR microswitches. We show that expansion of the orthosteric pocket is coupled to a distinct set of intracellular interactions that regulate the receptor’s elevated basal signaling. Our findings reveal a novel mechanism for ACKR3 activation that diverges from established models and illustrates the utility of nanobodies for targeting other “undruggable” GPCRs.

## Results

### The antagonist-bound ACKR3 structure adopts an active-like conformation

We previously developed a structural model for ACKR3 binding by the neutral antagonist nanobody VUN701^21^, and therefore solved the ACKR3–VUN701 complex by cryo-EM at a global resolution of 3.3 Å, expecting to capture an inactive receptor conformation. (Figure 1d, Supplementary Table 3, Supplementary Figure 1). To facilitate structure determination, VUN701 was engineered with three mutations near its C-terminus to permit binding of a nanobody-binding antibody fragment (NabFab)^22^ which was further stabilized by an anti-Fab nanobody^23^ to enlarge the system and facilitate particle picking (Supplementary Figure 2).

The solved structure aligned closely with our experimentally validated homology model of the VUN701-ACKR3 complex^21^, with an all-atom RMSD of 3.3 Å (Supplementary Figure 3). As predicted, density was resolved for residues C26–P35 of the ACKR3 N-terminus, which crosses the VUN701 β-sheet fold in an extended conformation (Figure 1e, left). The cryo-EM structure also clarified earlier uncertainties regarding the importance of CDR2 for VUN701 potency, revealing stable contacts between D56^CDR2^ and the backbone amides of C287^ECL3^ and R288^7×25^ (Figure 1e, right). The cryo-EM map also confirmed that VUN701’s H32^CDR1^ forms a cation–π interaction with F285^ECL3^ with stabilizing contacts from F106^CDR3^. Within the orthosteric pocket K100^CDR3^ engaged D275^6×58^ and E290^7×27^, while several CDR3 side chains made contacts with ECL2. Together, these features demonstrate that VUN701’s exceptional selectivity for ACKR3 is encoded by specific contacts with multiple elements of ACKR3’s extracellular surface.

In contrast, the intracellular face of ACKR3 appeared conformationally heterogeneous: regions of transmembrane helix 5 (TM5), TM6, TM7, and amphipathic helix 8 (H8) required for effector binding were unresolved, even after focused refinement (Supplementary Figure 1, Supplementary Figure 4). Three-dimensional variability analysis confirmed pronounced motion in TM6, TM7, and H8 (Supplementary Movie 1), consistent with ACKR3’s well-documented basal activity^24^. These observations suggest that the dataset captured a basal conformational ensemble in which ACKR3 samples both inactive and active-like states. Within this ensemble, only the structural features common to the basal state refined to high resolution, while dynamic intracellular regions remained unresolved. As a result, the intracellular faces of ACKR3 in the CXCL12- and VUN701-bound structures appear nearly indistinguishable (Figure 1f). Thus, even when bound to a neutral antagonist, ACKR3 continues to sample both inactive and active-like conformations, consistent with measurements in which VUN701 has no effect on ACKR3 levels at the plasma membrane, in contrast to inverse agonists that increase surface expression by suppressing basal arrestin recruitment^19^.

### ECL2 interactions control inverse agonism in ACKR3

We previously identified a single-residue variant of VUN701, T105A, that reduced receptor activity below basal levels while maintaining high-affinity binding, consistent with inverse agonist behavior^21^. Remarkably, other T105^CDR3^ substitutions yielded nanobody variants with distinct pharmacological properties, spanning inverse agonism to partial agonism (Figure 2a, Supplementary Table 4). Among these, VUN701 T105N displayed the strongest inverse agonism and was chosen as the best ligand option for stabilizing the inactive ACKR3 conformation.

**Figure 2.**
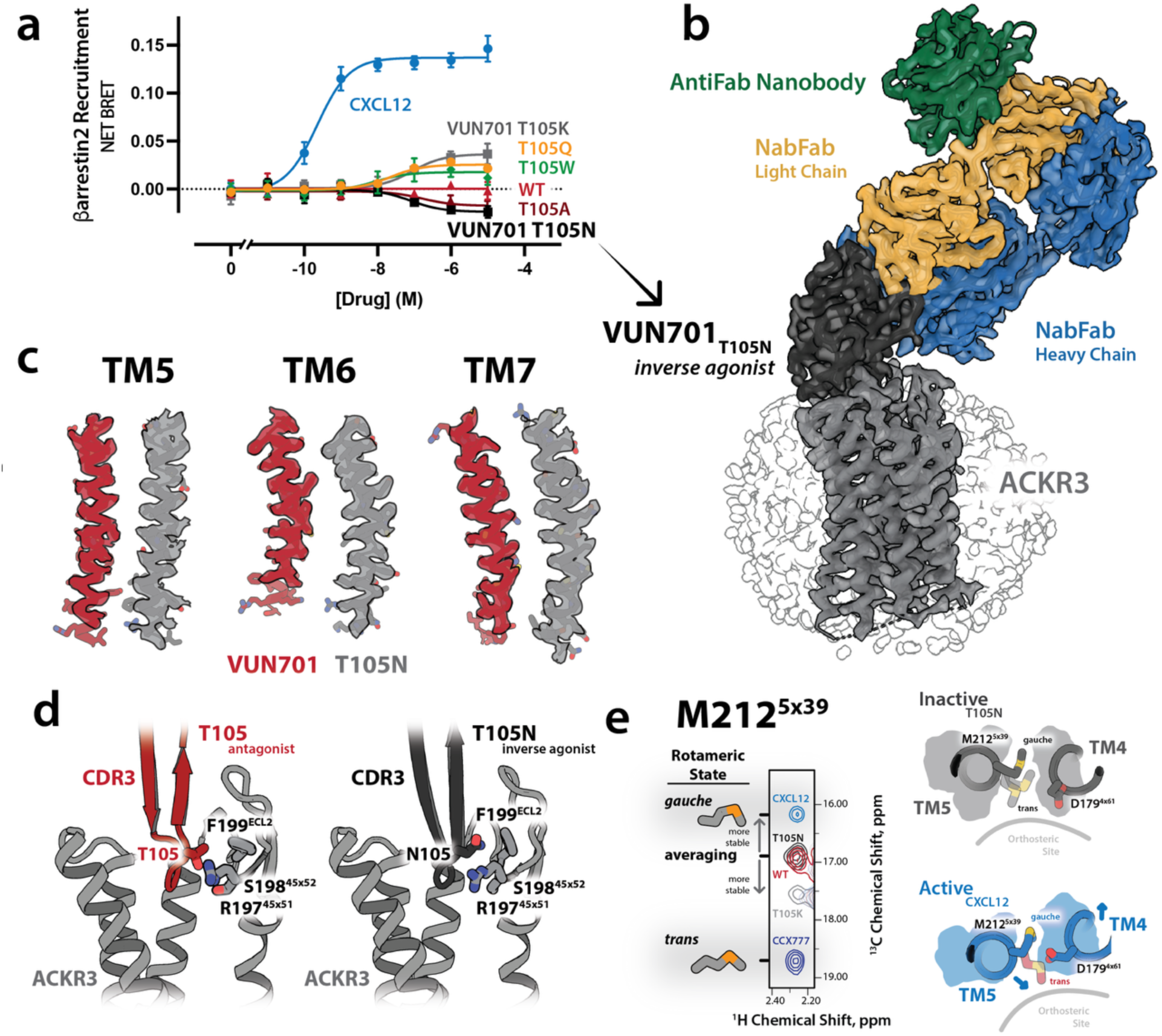
Extracellular Loop 2 (ECL2) movement drives ACKR3 inverse agonism. a) BRET-based β-arrestin2 recruitment assay for ACKR3 stimulated with the endogenous agonist CXCL12 (blue) or with VUN701 and VUN701 variants with single CDR3 substitutions at position T105. Mutations in T105 tune nanobody pharmacology from neutral antagonism (WT) to inverse agonism (T105A, T105N) and partial agonism (T105Q, T105W, T105K). b) Cryo-EM reconstruction of ACKR3 (gray) bound to the inverse-agonist nanobody VUN701 T105N (black). A nanobody-binding Fab (NabFab; light chain, orange; heavy chain, blue) and an anti-Fab nanobody (green) were included to increase particle mass and facilitate alignment. ACKR3 is shown embedded in an LMNG micelle (transparent surface). c) Comparison of transmembrane helices TM5, TM6, and TM7 between the antagonist VUN701–bound (red) and inverse agonist T105N-bound (gray) structures. The T105N complex stabilizes the intracellular face of TM5, TM6, and TM7, and has improved cryo-EM density. d) Structural comparison of the extracellular binding site highlights that VUN701 (red) and T105N (gray) adopt nearly identical overall binding poses, despite their opposing pharmacological effects. The T105N substitution alters ECL2 interactions. e) Left: ^1^H–^13^C NMR spectra of M212^5×39^ in ACKR3 bound to CXCL12, VUN701 WT, VUN701 T105N, VUN701 T105K, and CCX777 showing a transition from rotameric averaging in the inactive states to a stable rotameric state (gauche, trans) in the active states. Right: Structural views of TM4 and TM5 illustrating how movements in ECL2 lead to rearrangements in TM4 and TM5 reposition to restrict M212^5×39^ rotameric states. N = 3 biologically independent experiments plotted as mean +/- SEM.

The ACKR3–VUN701 T105N complex was solved by cryo-EM at 3.4 Å resolution (Figure 2b, Supplementary Table 3, Supplementary Figure 5). Although T105N binds in the same overall orientation as wild-type VUN701, it stabilizes a fully inactive receptor state with improved intracellular density and no detectable conformational heterogeneity in 3D variability analysis (Figure 2c). The side chain of N105 packs against ECL2 and makes no contacts with the orthosteric pocket or other microswitch motifs (Figure 2d). Since ECL2 is not regarded as important for activation of most class A GPCRs, the mechanism by which T105 substitutions affect ACKR3 activity was not readily apparent.

We previously established the side chain of M212^5×39^ as a sensitive NMR probe of receptor activation^18^. In agonist-bound active states, coordinated movements of TM4 outward and TM5 inward in response to ECL2 movement impose steric constraints on M212^5×39^, stabilizing a defined rotameric conformation (Figure 2e). For CXCL12, this corresponds to a gauche, outward-facing rotamer, whereas in the small-molecule agonist CCX777 the side chain adopts a trans, inward-facing orientation, likely due to larger transmembrane movements. In contrast, in the inverse agonist–bound T105N structure, M212^5×39^ is no longer sterically constrained, consistent with the rotameric averaging observed by NMR. The VUN701 T105K partial agonist variant produces an intermediate NMR signature between T105N and CCX777, indicating that this nanobody samples active-like conformations but does not fully stabilize the receptor (Supplementary Figure 6). Together, these observations provide a direct structural explanation for the M212^5×39^ NMR probe behavior and establish a mechanistic link between ECL2-driven extracellular remodeling, transmembrane helix rearrangements, and ACKR3 activation.

### Engineered nanobodies activate ACKR3 by displacing ECL2

Having established that VUN701 variants can activate ACKR3, we noted that bulkier T105^CDR3^ substitutions generally produced greater receptor activation (Figure 2a) and used ProteinMPNN^25^ and AlphaFold2^26^ to design five residue insertions in the CDR3 loop (Figure 3a). Each variant was predicted to fold with high-confidence and acted as an ACKR3 agonist, producing measurable β-arrestin recruitment in the absence of CXCL12 (Figure 3b, Supplementary Table 4). We solved the structure of ACKR3 bound to VUN-XL, the most efficacious agonist, by cryo-EM at 3.9 Å resolution in the presence of NabFab and CID24, a Fab selective for the fully active state of ACKR3^27^ (Figure 3c, Supplementary Table 3, Supplementary Figure 7). Like VUN701 and VUN701-T105N, VUN-XL bound near the extracellular surface, in contrast to the deep orthosteric engagement of CXCL12 and other GPCR agonists. In addition to the ACKR3-VUN701 contacts described above, VUN-XL formed new interactions with E212^5×40^ and W100^2×60^ (Figure 3d).

**Figure 3.**
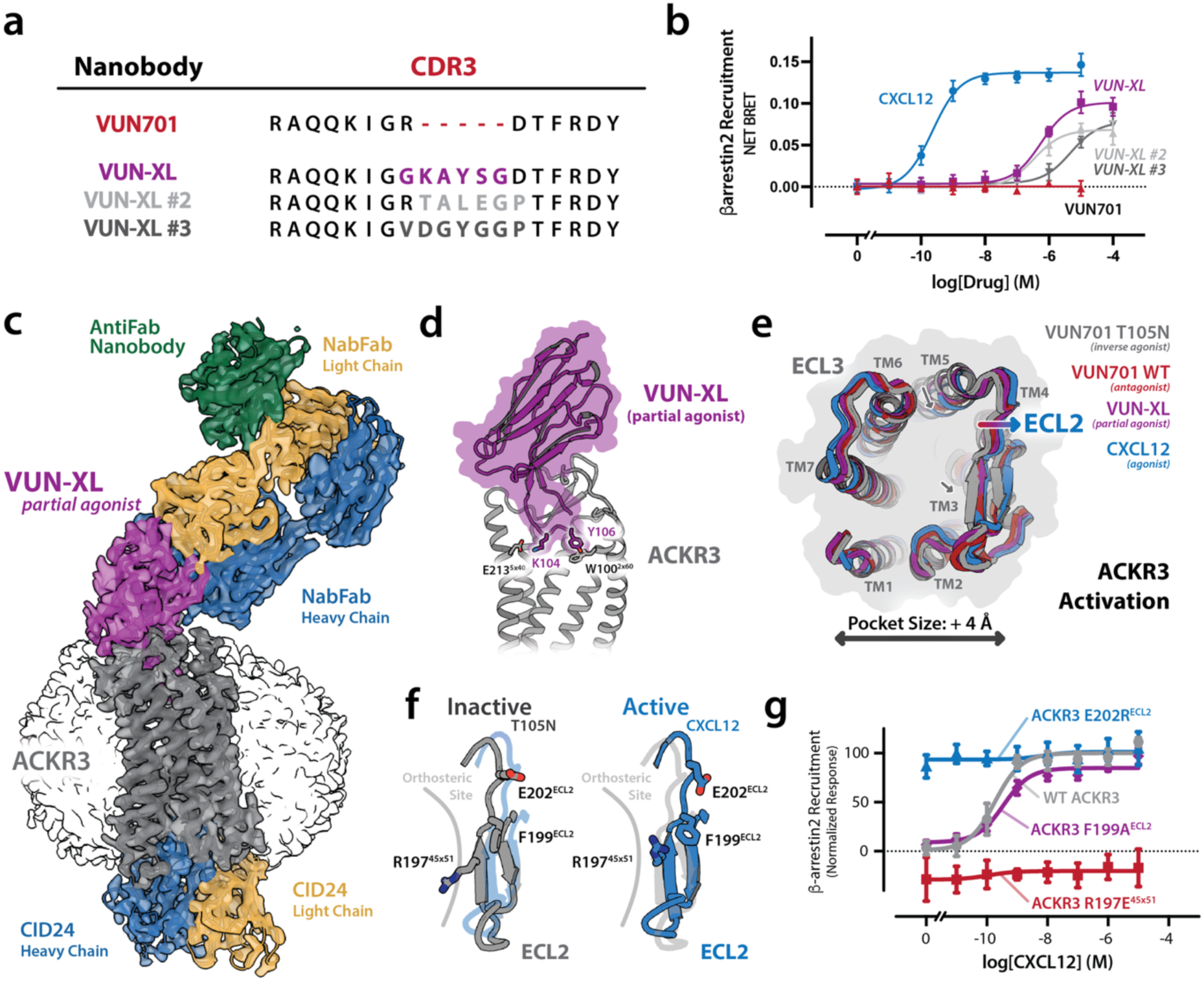
Orthosteric volume expansion governs activation of ACKR3. a) Alignment of CDR3 sequences for VUN701 and engineered CDR3-extended variants (VUN-XL, VUN-XL #2, VUN-XL #3). Inserted residues (highlighted) increase steric bulk within the extracellular vestibule while preserving the overall nanobody scaffold. b) BRET-based β-arrestin2 recruitment assay showing ACKR3 inhibition by VUN701 (red), and activation by CXCL12 (blue), and CDR3-extended nanobody variants. While VUN701 remains inactive, CDR3 extension confers agonist activity, with VUN-XL exhibiting the highest potency and efficacy. c) Cryo-EM reconstruction of ACKR3 (gray) bound to the agonist nanobody VUN-XL (purple). A nanobody-binding Fab (NabFab; light chain, gold; heavy chain, blue), an anti-Fab nanobody (green), and the active-state–selective Fab CID24 (light chain, gold; heavy chain, blue) were included to facilitate particle alignment and stabilize the active conformation. ACKR3 is embedded in an LMNG micelle (transparent surface). d) Close-up view of the VUN-XL–ACKR3 interface highlighting designed CDR3 residues K104 and Y105 engaging ACKR3 extracellular residues E212^5×40^ and W100^2×60^. e) Extracellular ribbon overlay of ACKR3 structures bound to VUN701 T105N (inverse agonist), VUN701 (antagonist), VUN-XL (partial agonist), and CXCL12 (agonist). Increasing receptor activity correlates with progressive outward displacement of ECL2 and coordinated rearrangements of TM4 and TM5, resulting in enlargement of the extracellular vestibule by >4 Å. f) Structural comparison of ECL2 in inactive (T105N) and active (CXCL12) states illustrates inward collapse versus outward opening of the loop, respectively, with accompanying repositioning of residues E202^ECL2^, F199^ECL2^, and R197^4×51^. g) β-arrestin2 recruitment for disruptive ACKR3 ECL2 mutants R197^45×51^ (red), F199^ECL2^ (purple), and E202^ECL2^ (blue) in response to CXCL12 monitored by BRET. Charge-reversal mutations in ECL2 (E202R, R197E) produce ligand-independent activation and inhibition, validating ECL2 as a dominant extracellular control element. N = 3 biologically independent experiments plotted as mean +/- SEM.

From an overlay of ACKR3 structures bound to VUN701 (neutral antagonist), T105N (inverse agonist), VUN-XL (partial agonist), and CXCL12 (full agonist), receptor activation appears proportional to ECL2 displacement and an overall widening by ∼4 Å of the opening to the orthosteric pocket (Figure 3e). The inactive VUN701-T105N complex exhibits an inward ECL2 position that occludes the orthosteric pocket, whereas the neutral antagonist and agonist nanobodies progressively expand this region. Outward displacement of ECL2 is accompanied by motions deeper in the transmembrane core: TM5 pivots inward toward the helical bundle while TM3 shifts outward along the extracellular face priming the intracellular side for activation.

To test the hypothesis that ECL2 displacement can activate the receptor, we mutated ACKR3 residues at or near the interface with CDR3 in the nanobody complexes (Figure 3f). Among the tested ECL2 substitutions, E202R^ECL2^ conferred strong ligand-independent activation, whereas R197E^45×51^ diminished basal signaling (Figure 3g, Supplementary Table 5). The dramatic reciprocal effects of these single residue substitutions paired with the 4 Å alteration in binding pocket diameter involving ECL2 supports the idea that ECL2 conformation is correlated with ACKR3 activation state. This reinforces the unexpected discovery that nanobodies can elicit the full pharmacological spectrum from inverse agonism to agonism simply by engaging ECL2.

### ACKR3 undergoes classical intracellular rearrangement through a conserved aromatic network

During ACKR3 activation, the inward movement of TM5 seen in the orthosteric site (Figure 3e) propagates through the transmembrane domain to the intracellular face of the receptor. In this inward state, Y232^5×58^ shifts toward the intracellular pocket to form a stabilizing interaction with Y315^7×53^, a pairing observed in other active-state GPCRs^28^. In ACKR3, this interaction is further reinforced by Y257^6×34^ and R142^3×50^ forming a four-residue cluster unique to this receptor (Figure 4a)^27, 29^. As an element of the conserved DRY motif, R142^3×50^ would ordinarily help stabilize the inactive receptor conformation^5^, but its engagement by the tyrosine triad of Y232^5×58^, Y315^7×53^ and Y257^6×34^ appears to increase the frequency with which ACKR3 adopts the active conformation^29^. Disrupting any of these four residues destabilized the network and reduced basal β-arrestin recruitment, comparable to the levels observed in a phosphorylation-deficient mutant (ST/A) (Figure 4b, Supplementary Table 6).

**Figure 4.**
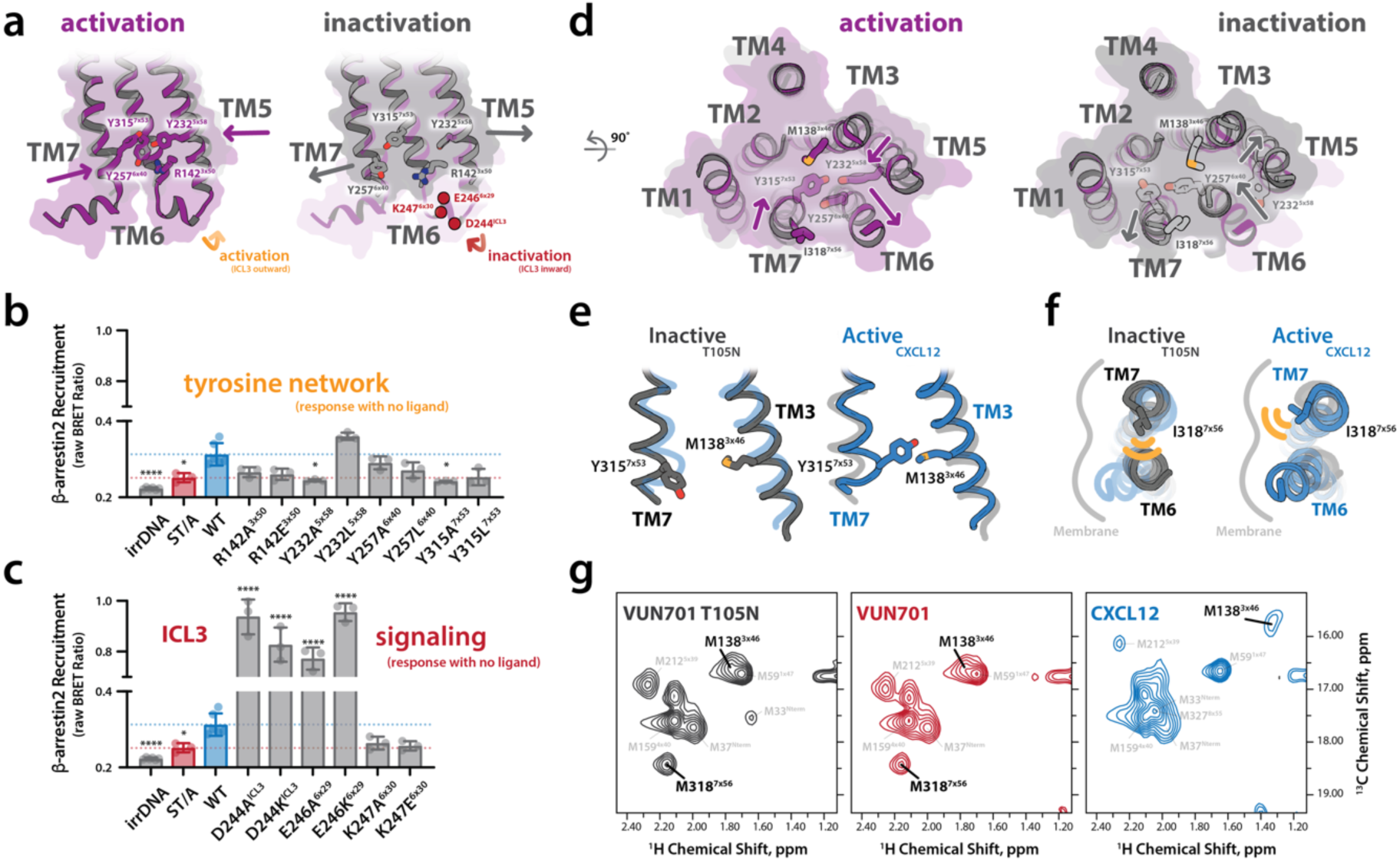
ACKR3 ligand-dependent activation is microswitch independent. a) Structural schematics comparing inactive (T105N) and active (VUN-XL) ACKR3 conformations highlight a conserved intracellular aromatic network that stabilizes receptor activation. In the active state (left, purple), inward movement of TM5 and TM7 promotes formation of a four-residue aromatic network involving Y232^5×58^, Y257^6×34^, Y315^7×53^, and R142^3×50^. In the inactive state (right, gray), this network is disrupted, and the intracellular helices adopt a relaxed configuration. b) Functional analysis of residues forming the intracellular aromatic network. β-arrestin2 recruitment measured by NanoBRET in the absence of ligand reveals that disruption of individual tyrosine residues within the network significantly reduces basal signaling, comparable to a phosphorylation-deficient mutant (ST/A). c) Functional interrogation of intracellular loop 3 (ICL3). Mutations of charged residues within ICL3 alter basal β-arrestin recruitment in a residue-specific manner, demonstrating that the accessibility of ICL3 acts as an active arrestin-interaction surface. d) Bottom-up intracellular views of ACKR3 illustrating global transmembrane rearrangements associated with activation (left, purple) and inactivation (right, gray). Despite its inability to couple heterotrimeric G proteins, ACKR3 undergoes canonical GPCR-like motions, including outward displacement of TM6 and inward movement of TM7 in the active state. The locations of methionine probes for activation and residues involved in the tyrosine network are highlighted. e) Structural comparison of TM7 between inactive (VUN701 T105N; gray) and active (CXCL12; blue) states reveals repositioning of TM7 and Y315^7×53^ toward the receptor core during activation. M138^3×46^ acts as a sensitive probe to sense the change in aromatic environment. f) TM6 and TM7 rearrangements accompanying ACKR3 inactivation (gray) and activation (blue). In the active state, TM6 swings outward relative to the membrane plane, consistent with a transducer-compatible conformation, whereas TM6 remains packed against the helical bundle in the inactive state. The mutation of I318^7×56^ to methionine creates a sensitive probe of TM6 movements. g) ^1^H–^13^C NMR spectra validating M138^3×46^ and I318M^7×56^ as reporters of TM7 and TM6 motion, respectively. In the inactive and basal states (VUN701, VUN701 T105N), M318^7×56^ peaks appear in a restrained rotameric state, consistent with a tightly packed TM6-TM7 environment. M138 is shifted away from the aromatic residue Y315^3×46^ on TM7. In the active state (CXCL12), spectral changes indicate an outward movement of TM6 with the disappearance of M318^7×56^, and an inward movement of TM7 with an aromatic shift in M138^3×46^. N = 3 biologically independent experiments plotted as mean +/- SEM.

This aromatic network not only stabilizes an ACKR3 active conformation but also appears to orchestrate the hallmark intracellular rearrangements of GPCR activation. Outward displacement of TM6 and inward motion of TM7—long considered structural signatures of G protein coupling—were both evident in ACKR3 despite its inability to engage heterotrimeric G proteins (Figure 4a).

In addition to expanding the intracellular pocket, these motions coincided with the exposure of charged residues on ICL3 (D244^ICL3^, E246^6×29^, and K247^6×30^), which appear essential for modulating arrestin recruitment. Mutation of any one of these residues either amplified or suppressed basal signaling, highlighting ICL3 as an active binding interface for β-arrestin rather than a passive loop (Figure 4c, Supplementary Table 6). These results align with recent work showing that arrestin can engage intracellular loop residues in certain GPCRs, including ACKR3^30^ bypassing the canonical G protein cavity, and suggest that ICL3 in ACKR3 has been repurposed as a specialized arrestin-binding platform.

Although ACKR3 is considered atypical because it does not couple to G proteins, its intracellular conformational dynamics mimic those of prototypical GPCRs (Figure 4d). Specifically, the outward shift of TM6 and inward shift of TM7 we observed are virtually indistinguishable from structural rearrangements that enable transducer binding in receptors like β_2_AR, A_2A_R, or rhodopsin. This paradox—ACKR3 structurally mimicking an active G protein–coupling state while functionally excluding G proteins—raises important questions about the kinetic and energetic origins of its signaling bias. To exclude artifacts from Fab scaffolding, we validated these conformational changes with independent NMR probes at ^13^C-ε-methyl-labeled M138^3×46^ and M318^7×53^ (a conservative substitution for I318). TM7 motion was confirmed by the strong upfield ring-current shift of M138^3×46^, a direct consequence of Y315^7×53^ moving inward (Figure 4e, Figure 4g); replacing Y315^7×53^ with leucine abolished this signal, providing direct evidence of its role (Supplementary Figure 8). Likewise, TM6 motion was confirmed by monitoring ^13^C-ε-methyl-labeled M318^7×56^: in the inactive state, M318^7×56^ produced a sharp peak consistent with packing against TM6, but in the active state, this signal broadened and collapsed into the random-coil region, consistent with TM6 swinging outward (Figure 4f, Figure 4g, Supplementary Figure 9).

ACKR3’s failure to engage G proteins despite adopting these conformations suggests that ACKR3’s signaling specificity is not encoded in its global architecture, but rather in amino acid-level changes in G protein contacts.

### ACKR3 can be activated by increasing orthosteric volume rather than classical microswitches

Our cryo-EM structures firmly establish that ACKR3, despite its atypical signaling profile, undergoes canonical transmembrane motions. Chemokines typically activate their receptors by penetrating the orthosteric pocket to engage conserved microswitches. In contrast, the binding of VUN701 ligands is mediated by elements at or near the extracellular surface, and the opening to ACKR3’s orthosteric pocket grows wider upon activation (Figure 3e). Volumetric analysis of all ACKR3 structures using POVME2 revealed a surprising correlation: pocket expansion accompanies receptor activation, whereas contraction coincides with inactivation (Figure 5a, Supplementary Figure 10, Supplementary Data 1). The inactive, T105N-bound structure exhibited the smallest pocket volume (536 Å^3^), while the other nanobody-bound complexes—VUN701 and VUN-XL— expanded to 857 and 947 Å^3^, respectively, despite VUN701 and T105N having exactly one amino acid difference and VUN701 and VUN-XL having only five additional residues.

**Figure 5.**
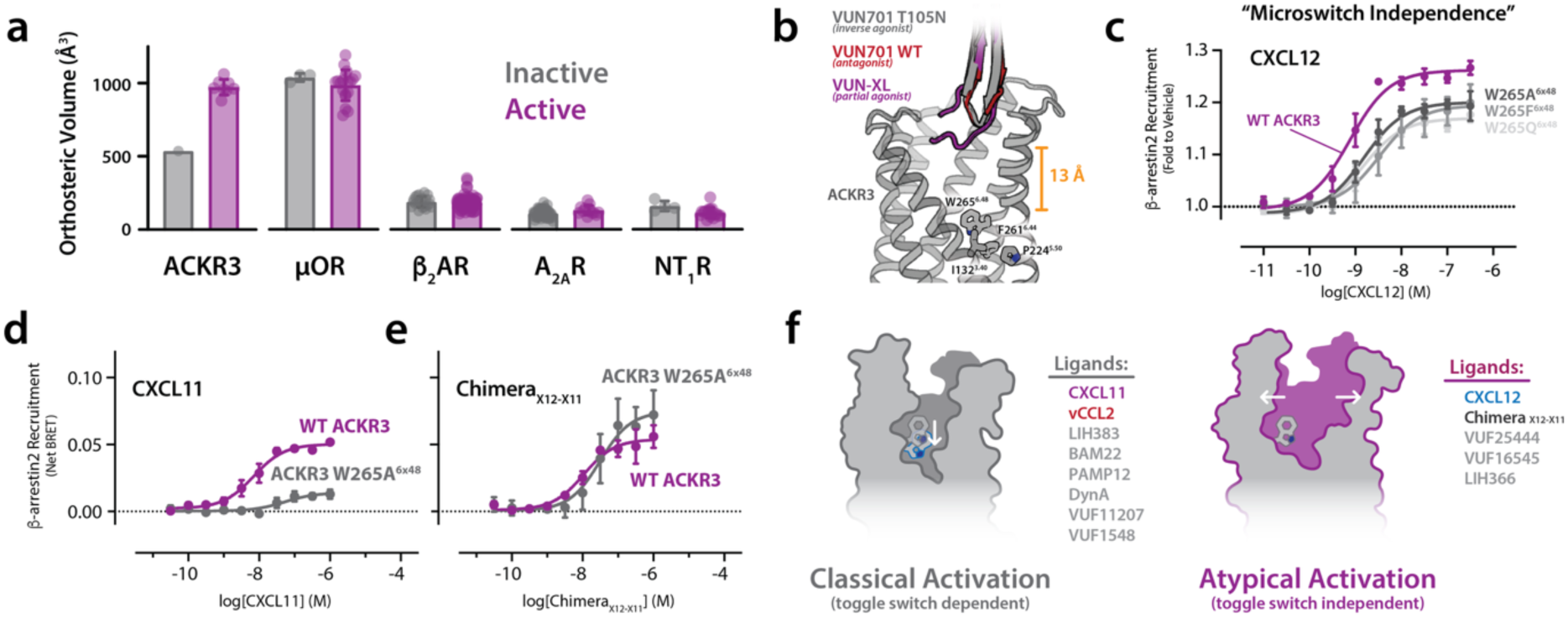
A unique aromatic network regulates ICL3 exposure and basal signaling. a) Orthosteric pocket volumes calculated for ACKR3 and representative class A GPCRs in inactive (gray) and active (purple) conformations. ACKR3 uniquely exhibits a large active-state expansion (∼500 Å^3^), whereas μOR, β_2_AR, A_2A_R, and NT_1_R show minimal change upon activation. b) Structural context for “microswitch independence.” Overlay of ACKR3 structures bound to VUN701 T105N (inverse agonist; gray), VUN701 WT (antagonist; red), and VUN-XL (partial agonist; purple) illustrating the shallow extracellular binding of all nanobodies. Conserved microswitch residues in the transmembrane core (including W265^6×48^, and the PIF region) are distal from the nanobody binding site; the closest approach to the toggle-switch region is >13 Å, consistent with modulation of activity without direct microswitch engagement. c) β-arrestin2 recruitment dose–response to CXCL12 for WT ACKR3 (purple) and toggle-switch mutants at W265^6×48^ (gray traces). Substitution of W265^6×48^ (A/F/Q) has minimal impact on CXCL12-driven arrestin recruitment, indicating that ACKR3 activation by CXCL12 does not require the canonical toggle-switch residue. d) β-arrestin2 recruitment dose–response to CXCL11 comparing WT ACKR3 (purple) to ACKR3 W265A^6×48^ (gray). In contrast to CXCL12, CXCL11-driven activation is strongly reduced by toggle-switch disruption, revealing ligand-dependent reliance on W265^6×48^. e) β-arrestin2 recruitment dose–response to a CXCL12–CXCL11 chimera (X12–X11) comparing WT ACKR3 (purple) to ACKR3 W265A^6×48^ (gray). The chimera displays toggle-switch independence akin to CXCL12. f) Working model summarizing two activation modes in ACKR3. Left: “Classical” GPCR activation in which ligand efficacy depends on the toggle switch (CXCL11, vCCL2, LIH383, BAM22, PAMP12, dynorphin A, VUF11207, VUF15485 – Supplementary Figure 11). Right: “Atypical” activation in which efficacy is toggle-switch independent and instead correlates with extracellular remodeling and orthosteric pocket expansion (CXCL12, Chimera_X11-X12_, VUF25444, VUF16545, LIH366). N = 3 biologically independent experiments plotted as mean +/- SEM.

This finding is highly unusual. In several studies of canonical GPCRs, the opposite relationship is described: activation is associated with contraction of the orthosteric pocket upon ligand engagement. This contraction arises from the packing of extracellular loops around the ligand to initiate conformational changes in conserved microswitches that propagate activation to the intracellular side^31,32^. Structural comparisons of inactive and active states and biophysical studies - such as pressure-induced activation of β_1_AR and β_2_AR - have established reduced pocket volume is a hallmark of the active conformation^33,34^. We confirmed this behavior by comparing pocket volumes across published GPCR structures, including MOR, A_2A_R, and NT_1_R (Supplementary Data 1). Only ACKR3 displayed a measurable increase in orthosteric volume upon activation (Figure 5a). This expansion likely arises from the ECL2-mediated rearrangement observed in our nanobody structures: outward displacement of ECL2 physically pushes TM3 and TM4 away from the receptor core, creating space for TM5 to pivot inward (Figure 3e).

Consistent with this idea of a unique activation mechanism, none of our nanobody-bound structures showed direct contact with the conserved microswitches that traditionally govern activation, including the PIF and toggle switch motifs (Figure 5b). All functional nanobody backbones were positioned more than 13 Å from these regions, indicating that they modulate activity without directly engaging side chains in the transmembrane core. To test whether these microswitches are dispensable for ACKR3 signaling, we mutated the highly conserved toggle switch residue W265^6×48^, a site essential for activation in prototypical class A GPCRs. In receptors such as β₂AR and CXCR4, substitutions at this position abolish signaling^35–38^. Surprisingly, alanine, phenylalanine, and glutamine substitutions at W265^6×48^ had little to no effect on CXCL12-induced β-arrestin recruitment to ACKR3 (Figure 5c, Supplementary Table 7). In contrast, ACKR3 activation by CXCL11 was W265^6×48^ microswitch-dependent (Figure 5d, Supplementary Figure 11). Because a chimeric CXCL12 containing the CXCL11 N-terminus^39^ was unaffected by the toggle switch mutation (Figure 5e), we concluded that the CXCL12 N-terminus is dispensable and its well-documented interactions with ECL2^27,40,41^ are sufficient to induce β-arrestin recruitment.

Taken together, these findings revealed that ACKR3 can be activated in a W^6×48^ toggle switch-independent manner. Structural comparisons indicate that outward displacement of ECL2 (Figure 3e) expands the orthosteric pocket (Figure 5a) and drives conformational coupling through TM3, TM4, and TM5 to an intracellular aromatic network (Figure 4d and e). This mechanism may explain how diverse ligands—including opioids, nanobodies, chemokines, and cytokines—can all activate ACKR3 without deep penetration of the orthosteric pocket. To test this, we measured the impact of the W265A^6×48^ mutation on the signaling of a large panel of ACKR3 agonists. Ligands that activate ACKR3 through a toggle switch-dependent or -independent mechanism formed two groups, each of which contained small molecule, peptide and protein agonists (Figure 5f, Supplementary Table 8, Supplementary Figure 11). Collectively, these results challenge the long-held assumption that pocket contraction and microswitch toggling are universal hallmarks of GPCR activation and may help explain why ACKR3 inhibitor development is challenging.

## Discussion

ACKR3 has attracted significant attention among chemokine receptors and GPCRs due to its atypical signaling properties, ligand promiscuity, and proposed role in analgesia as a scavenger of endogenous opioid peptides^9,42,43^. Despite this intense interest, the inactive conformation of ACKR3 has remained unresolved, leaving a critical gap in understanding its structural dynamics. In this study, we used an inverse agonist nanobody to obtain the first bona fide inactive ACKR3 structure by cryo-EM for comparison with active-state structures bound to closely related ligands. Because the neutral antagonist VUN701 binds the receptor in a basally active ensemble of states, engineered nanobodies were crucial to revealing the molecular details of ACKR3 activation.

Uncovering this activation mechanism relied on precise nanobody engineering. Although nanobodies are widely used for GPCR structural studies, their application as pharmacological probes is limited^44^. We show that, with structural insight, nanobodies that bind the GPCR orthosteric pocket can be easily adapted as selective tools for probing and modulating receptor function. Because VUN701 and its variants share identical scaffolds and vary only in their CDR3 sequences, the effects of agonism, antagonism, and inverse agonism could be ascribed to specific ACKR3 conformational changes that might be less apparent if they were achieved with structurally diverse ligands. This underscores the value of nanobodies as tools for research and drug development.

ACKR3 is the most well-studied of the intrinsically β-arrestin biased atypical chemokine receptors, which engage β-arrestin in response to ligand binding but do not evoke G protein signaling. Among the atypical subfamily, ACKR3 responds to an unusually diverse set of protein, peptide and small molecule ligands (Figure 1a and b) that includes enkephalin and dynorphin peptides and other endogenous opioid receptor ligands^9^. Owing to its basal internalization and recycling activity, ACKR3 functions as a scavenger for the chemokines CXCL11 and CXCL12, and internalizes endogenous opioid peptides in the brain, regulating their availability for classical opioid receptors. ACKR3 scavenging of CXCL12 was recently exploited for targeted degradation of extracellular proteins^45^. Most chemokine receptors recognize their ligands based on a precise N-terminal sequence that makes specific contacts deep in the binding site. However, it was previously observed that ACKR3 efficiently scavenges the N-terminally processed chemokines that no longer activate their conventional receptors CXCR3 and CXCR4^12,46^. Thus, ACKR3’s broad ligand binding profile and insensitivity to N-terminal ligand truncation hinted at a divergent activation mechanism for the atypical receptor.

Our structure-function analysis of ACKR3 reveals a new mechanism that challenges the canonical paradigm of GPCR activation. As an alternative to engagement of conserved microswitches at the base of the orthosteric pocket, the atypical activation of ACKR3 employs extracellular loop remodeling and a cascade of conformational changes culminating in intracellular network assembly. Single mutations in ECL2 that prevent ACKR3 activation or lock it into a constitutively active conformation (Figure 3f) reinforce our observation that ACKR3 can be activated independently of the W265^6×48^ toggle switch. Activation by enlargement of the orthosteric pocket (Figure 5f) explains ACKR3’s promiscuous scavenging of diverse sets of ligand scaffolds, high propensity for activation, and why ACKR3 inhibitor development has been historically challenging. Moreover, this alternative mechanism challenges the paradigm in which conserved microswitches are indispensable to GPCR function. ACKR3’s ECL2-triggered activation mechanism is likely shared with other chemokine receptors and may be relevant to other GPCRs. In structure-function studies of CCR6 andCCR9, activation by chemokine ligands with truncated N-terminal sequences occurs independent of deep orthosteric contacts^47,48^. In particular, CCR9 activation tolerates alanine mutations in the CCL25 N-terminus but is strongly affected by modifications to the 30s loop, which is predicted by homology modeling to make specific contacts with ECL2^47^. Other receptors, like ACKR1 and ACKR5, appear to have evolved fixed conformational states, completely eliminating transducer binding or enabling constitutive transducer interactions, respectively^49–53^. Given their close phylogenetic relationship, ACKR1 and ACKR5 may share ACKR3’s toggle switch-independence while acquiring mutations in ECL2 or in other regions that lock the extracellular pocket closed or open.

In most GPCRs, the conserved DRY motif in TM3 stabilizes the inactive state^5^, but ACKR3 contains an unusual aromatic cluster formed by Y232^5×58^, Y257^6×34^, Y315^7×53^, and R142^3×50^. This dynamic network at the intracellular pocket (Figure 4a) seems to control basal activity of ACKR3 by destabilizing canonical DRY contacts. Mutational disruption of the aromatic cluster restores DRY stabilization of the inactive state and reduces basal β-arrestin recruitment (Figure 4b), offering a clear structural rationale for ACKR3’s basal activity. Beyond the chemokine family, dozens of orphan GPCRs and the 70% of undrugged receptors may rely on similar noncanonical architectures, including receptors previously dismissed as “undruggable”, like ACKR3.

By elucidating how changes in the orthosteric pocket propagate to the ACKR3 effector binding site, we revealed a novel mode of GPCR signal transduction that deviates from the canonical microswitch paradigm. While ACKR3 employs some hallmarks of GPCR activation, like the outward movement of TM6 that opens the intracellular pocket, details revealed by these cryo-EM structures and mutational studies suggest that GPCRs cannot be fully understood through a single mechanistic lens. These findings highlight an important limitation in prevailing GPCR models: despite the fact that the majority (70%) of GPCRs remain untargeted, most therapeutic strategies continue to rely on mechanistic assumptions derived from a small subset of well-studied receptors. Notably, recent large-scale analyses indicate that many orphan GPCRs possess atypical extracellular loop 2 (ECL2) architectures and exhibit elevated basal or constitutive activity, raising the possibility that noncanonical, ECL2-driven activation mechanisms may be more widespread than previously appreciated and could contribute to the long-standing difficulty in pharmacologically targeting these receptors^54^. Our results suggest that many currently “undruggable” GPCRs may evade modulation not because they lack pharmacological potential, but because their activation mechanisms fall outside classical paradigms. Broadening our view of GPCR activation to include alternative allosteric pathways, such as extracellular loop–driven pocket expansion, may therefore enable continued GPCR drug development on previously inaccessible targets, expanding the therapeutic reach of the GPCR superfamily beyond its current boundaries.

## Methods

### Nanobody Expression and Purification

The DNA sequence encoding VUN701 and the AntiFab Nanobody were codon-optimized for E. coli expression and ordered from GenScript. Nanobodies were sub-cloned into a pET28a-6xHis-SUMO3 vector and expressed in BL21 DE3 E.coli. Single amino acid substitutions were made using Quikchange mutagenesis (Product Number: 200524 – Agilent), while designed extensions were made using Q5 Site-Directed mutagenesis (Product Number: E0554 – New England Biolabs). Cells were expressed at 37°C in Luria-Bertani (LB) medium and induced with 1 mM isopropyl-β-D-thiogalactopyranoside (IPTG) at an OD600 of 0.8. Cultures continued to grow for ∼6 hours before bacteria were pelleted by centrifugation and stored at −20°C. Bacterial pellets were resuspended in ∼20 mL of Buffer A (50 mM Na_3_PO_4_ (pH 8.0), 300 mM NaCl, 10 mM imidazole, 1 mM phenylmethylsulphonyl fluoride (PMSF), and 0.1% (v/v) 2-mercaptoethanol (BME)) per pellet and lysed via sonication. Lysed cells were clarified at 18,000 x g and the supernatant was discarded. Pellets were resuspended by sonication in ∼20 mL of Buffer AD (6 M guanidine hydrochloride, 50 mM Na_3_PO_4_ (pH 8.0), 300 mM NaCl, 10 mM imidazole) and spun down at 18,000 x g for 20 min. Using an AKTA-Start system (GE Healthcare), supernatant was loaded onto a Ni-NTA column equilibrated in Buffer AD. The column was washed with Buffer AD, and proteins were eluted using Buffer BD (6 M guanidinium, 50 mM sodium acetate (pH 4.5), 300 mM NaCl, and 10 mM imidazole). Proteins were refolded overnight via drop-wise dilution into a 10-fold greater volume of Refold Buffer (50 mM Tris (pH 7.6), 150 mM NaCl) with the addition of 35 mM cysteine, and 0.5 mM cystine. Refolded protein was concentrated in an Amicon Stirred Cell concentrator (Millipore Sigma) using a 10 kDa membrane. Concentrated protein was added to 6-8 kDa dialysis tubing with the addition of ULP1 to cleave the N-terminal 6xHis-SUMO3-tag and dialyzed at 25°C against Refold Buffer overnight. The AKTA-Start system was used to load the cleaved protein onto a Ni-NTA column equilibrated in Nanobody Buffer A (Refold Buffer + 10 mM Imidazole). The column was washed with Nanobody Buffer A, and the protein was eluted using Nanobody Buffer B (Refold Buffer + 500 mM Imidazole). Each nanobody underwent four rounds of dialysis in 5 mM ammonium bicarbonate, lyophilized, and stored at −80°C for further use. Purity and identity of nanobodies was confirmed by electrospray ionization mass spectrometry using a Thermo LTQ instrument and SDS-PAGE with Coomassie staining and found to be >99% pure.

### Chemokine Purification

All chemokines utilized in these experiments, including CXCL12, CXCL11, vCCL2, and Chimera_X12-X1139_ were purchased from Protein Foundry, L.L.C.

### NabFab Expression and Purification

A pRH2.2 duet plasmid encoding the NabFab construct was expressed and purified following previously described procedures with minor modifications. Briefly, BL21(DE3) cells harboring the plasmid were grown in autoinduction medium (Terrific Broth supplemented with 0.02% lactose and 0.01 % glucose) at 37 °C for 6 h, followed by overnight expression at 18 °C. Cells were harvested by centrifugation and stored at –20 °C until purification.

Frozen pellets were resuspended in Buffer A (50 mM Tris pH 8.0, 150 mM NaCl, 1 mM PMSF) at ∼20 mL per liter of culture and lysed by sonication. Lysates were incubated at 60 °C for 30 min to enrich thermally stable Fab fragments and then clarified by centrifugation at 18,000 × g. The supernatant was applied to a Protein L affinity column (Cytivia Product 17547815), washed with Buffer A, and eluted with Acetic Acid, pH 3). The pooled elution containing NabFab was immediately neutralized, pooled, and concentrated.

Purified NabFab was estimated to be >95% pure by SDS–PAGE and was used directly for downstream complex assembly.

### ACKR3 Expression

Recombinant baculovirus was produced using the Bac-to-Bac Baculovirus Expression System (Invitrogen). A pFastBac1 plasmid containing the full-length ACKR3 construct containing a C-terminal, 3C cleavable FLAG and 10xHis tag was transformed into DH10Bac E. coli (Thermo Fisher) and subsequently plated onto LB agar with 50 μg /mL, kanamycin, 7 μg /mL gentamicin, 10 μg /mL tetracycline, 100 μg/mL Bluo-Gal and 40 μg /mL, IPTG (Teknova). Blue/white screening identified recombinant (white) colonies, of which individual clones were inoculated in 5mL of LB with 50 μg /mL, kanamycin, 7 μg /mL gentamicin, and 10 μg /mL tetracycline and placed in a 37°C shaking incubator overnight. Cultures were pelleted, lysed, and neutralized using buffers from the GeneJET Plasmid Midiprep Kit (Thermo Fisher Scientific) and bacmid was purified by isopropanol precipitation. Final bacmid pellets were solubilized in 40 mM Tris, pH 8 and 1 mM EDTA. Sf9 cell transfection used a mixture of purified bacmid (5μl – 1µg total DNA), X-TremeGENE HP DNA (3 μl) and Expression Systems Transfection Medium (100 μl) was mixed and added to 2.5 ml of Sf9 cells at ∼ 1.2 × 106 cells/mL and the bacmid-cell mixture was placed in a 24-well, deep-well plate (Thomson Instrument Company) covered with a polyurethane sealing film (Diversified Biotech). Cells were incubated at 27 °C for 96 hours at 300 rpm. Cells were subsequently pelleted, and the supernatant was collected to isolate P0 virus. P0 virus titers were determined using gp64 titer assay^55^. P1 (“first passage”) virus was produced by infecting 50 ml of Sf9 cells at a density of ∼ 2.0 x 10^6^ cells/mL with titered P0 virus at an MOI of 0.1-0.5. Cells were incubated at 27 °C for 72 hrs shaking at 144rpm. Cells were pelleted and the supernatant was collected and titered using the same gp64 titering assay^55^. Large-scale expression of labeled ACKR3 was performed by adding high titer P1 virus (≥ 1 x 10-9 IU/mL to ∼2 L of Sf9 cells in normal media (cryo-EM) or methionine-deficient medium (NMR - Expression Systems) at a density of 3.5-4.0 x 10^6^ cells/mL at an MOI of 5. Methionine-labeled ACKR3 for NMR experiments was produced by adding 250 mg L-1 13CH3-methionine (Cambridge Isotope Laboratories) cells 5 hours post-infection. Cells were incubated at 27 °C for 48 hours, then pelleted and stored at −80°C.

### ACKR3 Purification

Frozen cell pellets were diluted 1:1 in hypotonic lysis buffer (10 mM HEPES pH 7.5, 20 mM KCl, 10 mM MgCl_2_, Roche Complete Protease Inhibitor Cocktail, 2 mg/mL iodoacetamide) and thawed on ice. Direct solubilization was performed by passing the cell slurry through a 16-gauge needle four times to aid solubilization by lysing cells. Cell slurry was added to solubilization buffer (100 mM HEPES pH 7.5, 800 mM NaCl, 1.5% / 0.3% LMNG (Lauryl Maltose Neopentyl Glycol) /CHS (Cholesteryl Hemisuccinate)) and incubated at 4°C for 4 hours with stirring. The mixture was spun down at 50,000 x g for 30 min and the supernatant (∼200 ml) was transferred to 4x 50 ml conical tubes. 4 ml TALON cobalt resin slurry (Takara Bio Inc.) was added (1 ml/tube) with 10mM imidazole final concentration to limit non-specific binding, and the supernatant mixture was rocked overnight at 4°C. This mixture was added to columns the following day, and cobalt resin was washed with 20 ml of two wash buffers (Wash Buffer 1: 50mM HEPES pH 7.5, 400mM NaCl, 0.1%/0.02% LMNG/CHS, 10% glycerol, 20mM imidazole; Wash Buffer 2: 50mM HEPES pH 7.5, 400mM NaCl, 0.025%/0.005% LMNG/CHS, 10% glycerol, 10mM imidazole). Receptors were eluted with a high imidazole buffer (50mM HEPES pH 7.5, 400mM NaCl, 0.025%/0.005% LMNG/CHS, 10% glycerol, 250mM imidazole. Elutions were concentrated to 500 µL using a 30,000 MCWCO Amicon Ultra-4 Centrifugal Filter Unit (Millipore Sigma) and buffer exchanged into Exchange Buffer (25mM HEPES pH 7.5, 150mM NaCl, and 0.025%/0.005% LMNG/CHS) using a PD-10 desalting column (GE). Prescission Protease and PNGaseF were added to purified receptors overnight. The next day, TALON cobalt resin was added, and the mixture was incubated with rocking for 2 hours at 4°C. The mixture was added to a new column to separate cleaved receptor from the tag-bound cobalt resin and washed with Exchange Buffer to collect the flow through. Flow through was concentrated to 0.5 mL before size-exclusion chromatography (SEC) on a S200 10/300 Increase column (Cytiva), followed by further concentration and quantification.

### Assembly of Cryo-EM Complexes

Protein complexes were assembled by sequential addition of binding proteins followed by SEC. For VUN701 and VUN-T105N preparations, ACKR3 was first incubated with a 1:1.5 molar ratio of NabFab for 45 minutes at 4 °C. This incubation was followed by the addition of a 1:2 molar ratio of the anti-Fab nanobody. The mixture was again incubated for 45 min at 4 °C to allow complex formation prior to SEC purification.

For CID24 samples, NabFab and CID24 were added simultaneously to ACKR3 at a 1:1.5 molar ratio, after which the anti-Fab nanobody was added at a 1:3 molar ratio and the complex was incubated for 45 min at 4 °C. All complexes were purified by SEC to isolate the fully assembled species for cryo-EM preparation.

### Vitrification and Clipping (VUN701)

2.75 µl purified complex at a concentration of 7.5 mg/mL was applied to UltrAuFoil 1.2/1.3 300 mesh gold support films (Quantifoil), which had been glow discharged for 90 s. Samples were vitrified by plunge-freezing into liquid ethane using the Vitrobot Mark IV System (Thermo Fisher Scientific) while maintaining 100% humidity at 4 °C in the blotting chamber, and using a blot time of either 1.5 s or 3.0 s, a blot force of 0, and a wait time of 10 s. Support films were clipped using AutoGrid carrier assemblies (Thermo Fisher), immediately prior to imaging.

### Cryo-EM Data Collection (VUN701)

10,906 movies of ACKR3-VUN701-NabFab-AntiFabNanobody embedded in ice were recorded using a Krios G3i 300 kV microscope (Thermo Fisher) equipped with a K3 direct detection camera (Gatan) and a BioQuantum K3 imaging filter (Gatan). A nominal magnification of 105,000 x was used, with a physical pixel size of 0.85 Å/pixel. Movies were acquired as 60 dose fractions with a total dose of 50.3 e-/ Å^2^, yielding 0.833 e-/ Å^2^/frame. Fringe-free (aberration-free) imaging enabled a 3 x 3 multi-shot pattern and two shots per hole.

Samples were screened on a 200 kV Glacios microscope (Thermo Fischer) using a magnification of 54,000x, a physical pixel size of 0.73 pixel/Å. Each movie consisted of 100 frames collected with a total dose of 60.0 e-/Å^2^ giving 0.6 e-/ Å^2^/frame. Screening data was used to generate a high-quality template for particle picking.

### Vitrification and Clipping (VUN701 T105N/VUN-XL)

Immediately prior to vitrification, 8 mM CHAPSO (3-[(3-Cholamidopropyl) dimethylammonio]-2-hydroxy-1-propanesulfonate) was added to the purified complexes. 3 µL purified complex at a concentration of 7.5 mg/mL was then applied to C-flat 1.2/1.3 300 mesh copper support films (EMS), which had been glow discharged for 90 s. Samples were vitrified by plunge-freezing into liquid ethane using the Vitrobot Mark IV System (Thermo Fisher) while maintaining 100 % humidity at 4 °C in the blotting chamber, and using a blot time of 7.5 s, a blot force of −1, and a wait time of 3 s. Support films were clipped using AutoGrid carrier assemblies (Thermo Fisher).

### Cryo-EM Data Collection (VUN701 T105N/VUN-XL)

19,491 movies and 9,491 movies were recorded of the VUN701 T105N and VUN-XL Cryo-EM complexes, respectively using a Glacios2 200 kV microscope (Thermo Fisher) equipped with a Falcon4 direct detection camera (Thermo Fisher) and a Selectris imaging filter (Thermo Fisher). A nominal magnification of 165,000x was used, with a physical pixel size of 0.71 Å/pixel. Movies were acquired as 40 dose fractions with a total dose of 60 e-/Å^2^, yielding 1.5 e-/Å^2^/frame.

### Cryo-EM Data Processing

Dose-weighted micrographs were imported into cryoSPARC v4.5.1 (Structura Biotechnology). Motion correction and contrast transfer functions were calculated using the Patch Motion Correction and Patch CTF Estimation tools, respectively. Using the Curate Exposures tool, low-quality micrographs were manually rejected. Particles were extracted with a box size of 480 pixels binned to 120 pixels and 2D Class Averages were generated to yield a preliminary ab initio model. Fifty 2D class averages were created from the ab initio model and used as templates to re-pick particles, with a minimum separation distance of 0.3. The resulting particles were sorted with the Heterogeneous Refinement tool into the ab initio volume or into three scrambled volumes obtained by terminating the Ab-Initio Reconstruction tool before the first iteration. Particles sorted into the correct class underwent successive rounds of Heterogeneous Refinement before being re-extracted with a box size of 240 and then 480. Final models were created from full-resolution particles that underwent Non-Uniform Refinement, Local Refinement, and Reference-Based Motion Correction.

### Model Building and Refinement

Model building and refinement were carried out using an Alphafold2 predicted structure as a starting model, which was fitted into the map using UCSF ChimeraX. An initial model was generated using ISOLDE^56^. Models were subjected to multiple rounds of manual adjustment in COOT (v0.9.8.92) followed by automated real-space refinement using Phenix v1.21.5419^57^.

### ACKR3 Nuclear Magnetic Resonance (NMR)

Purified protein samples concentrated to ∼350 µL with 10% D_2_O by volume were loaded into a 5mm Shigemi microtube. ^13^C-^1^H HSQC spectra were collected at 310 K on a Bruker Avance Neo 800 MHz spectrometer equipped with a triple-resonance cryogenic probe. Each FID was the average of 512 transients with a time domain of 1024 points. The ^13^C dimension was composed of 90 complex points. Each scan was 8.9 minutes, with a total experimental time of 26 hours and 45 minutes. Data were processed using NMRPipe^58^ and visualized in XEASY^59^.

### Bioluminescence Energy Resonance Transfer (BRET) β-Arrestin Recruitment Assays

BRET experiments were performed to provide a direct and high-throughput readout of receptor activity in response to functional ligand mutants. The full-length sequence of ACKR3 was cloned into a pcDNA3.1 vector to include an N-terminal HA-FLAG tag and a C-terminal IDTG linker preceding the Rluc8 gene. Amino acid substitutions were made using Quikchange mutagenesis (Product Number: 200524 – Agilent). A pcDNA3.1 vector containing β-arrestin2 with an N-terminal Venus tag was also used in this assay. HEK293T cells (ATCC - CRL-3216) maintained in Dulbecco’s modified Eagle’s medium (DMEM) supplemented with 10% fetal bovine serum (FBS) were transiently transfected with 0.3 µg of receptor-Rluc8 and 5 µg of Venus β-arrestin at ∼50% confluency using TransIT-293 (Mirus). At 24 hours post-transfection, cells were resuspended in PBS supplemented with 0.1% glucose and plated in 96 wells plates at a density of 100,000 cells/well in a total of 60 µL. After an additional 1 hour at 37 °C, cells were stimulated with Coelenterazine-H to a final concentration of 5 µM and incubated for 5 minutes. Ligands suspended in PBS supplemented with 0.1% glucose were added at concentrations from 10 pM to 100 µM. WT-ACKR3:CXCL12 was included in each plate as a positive control and BRET (540/480 nm) signals were measured after 10 minutes on the Mithras LB940 (Berthold Technologies) using MicroWIN2010 5.19. Measurements on individual plates were performed in technical duplicates for each ligand concentration. Data were analyzed with a fixed slope (Hill Slope = 1), nonlinear fit to create a concentration-response curve in GraphPad Prism 9 or 10 (Graphpad Software Inc., San Diego, CA). Data from three biologically independent plates were used in the analysis and calculation of standard error. All data were baseline corrected to wells collected in the absence of functional ligands.

### NanoBRET-Based β-Arrestin Recruitment Assays

Constitutive and ligand-induced β-arrestin recruitment was monitored by NanoBRET as described previously^60^.

#### Constitutive β-arrestin recruitment

1.8 x 10^6^ HEK293T cells were seeded in 6-well plates and 24 hours later transfected with a vectors encoding ACKR3 variants C-terminally tagged with mNeonGreen and a vector encoding β-arrestin2 N-terminally tagged with Nanoluciferase. 24 hours after transfection, cells were resuspended in Opti-MEM and seeded in 96-well plates at a density of 1 x 10^5^ cells/well. Cells were incubated with coelentrazine-H (10 μM) for 15 minutes at 37°C. NanoBRET was immediately measured with a GloMax Discover plate reader (Promega) equipped with 450/8-nm filter for donor luminescence emission and 530-nm LP filter for acceptor fluorescence emission.

Measurements were performed in technical duplicates for each variant. Data points represent the mean of at least three independent experiments and are shown as raw NanoBRET ratios. Data analysis and plotting were performed using GraphPad Prism version 10.6.1.

#### Ligand-induced β-arrestin recruitment

5 x 10^6^ HEK293T cells were seeded in 10-cm dishes and 24 hours later transfected with vectors encoding ACKR3 variants C-terminally tagged with Nanoluciferase and a vector encoding β-arrestin2 N-terminally tagged with mNeonGreen. 24 hours after transfection, cells were resuspended in Opti-MEM and seeded in 96-well plates at a density of 1 x 10^5^ cells/well. Cells were incubated with coelentrazine-H (10 μM) for 15 minutes at 37°C. Ligands diluted in Opti-MEM at indicated concentrations were then added and NanoBRET was measured with a GloMax Discover plate reader (Promega) equipped with 450/8 nm filter for donor luminescence emission and 530-nm LP filter for acceptor fluorescence emission. Data were corrected to baseline signal measured in in the absence of ligands. Concentration–response curves were generated from the mean of at least three independent experiments using a three-parameter Hill equation in GraphPad Prism version 10.6.1.

### Computational Nanobody Design (VUN-XL)

The VUN701 sequence underwent multiple rounds of successive AlphaFold 2 modeling to create a longer CDR3 that maintained a similar structure. In the end, five residues (TALEGP) were chosen to replace D104, extending VUN701’s CDR3 architecture. Once the VUN701 CDR3 length and structure were optimized, the model of VUN701 with this long CDR3 was superimposed onto WT VUN701 in the ACKR3-VUN701 structure. This hybrid model was used as a structural input for ProteinMPNN^25^. Only the terminal CDR3 residues (KIGTALEGPTF) were allowed to be altered during the course of the computational ProteinMPNN run. Cysteines were excluded from the list of available residues to install. Using a temperature of 0.1, 100 sequences were designed onto the long-VUN701 backbone. Following the completion of the ProteinMPNN run, a sequence logo was generated. The top-scoring sequence, as well as the residues most utilized at each position in the sequence logo were used to create two new VUN701 protein sequences. AlphaFold models were generated to confirm the interactions of the longer-VUN701 designs and ACKR3 within the anticipated binding site. Each new nanobody was then cloned, purified, and tested in a BRET-based ACKR3 β-arrestin2 recruitment assay to determine potency and efficacy.

One of the two ProteinMPNN-generated nanobodies (design 1) displayed weak binding and efficacy in a BRET-based ACKR3 b-arrestin2 recruitment assay. The other generated nanobody no longer bound ACKR3. The computational model of design 1 was then used as a template to repeat manual nanobody generation and sequence optimization with ProteinMPNN. After 2 additional rounds of successive nanobody design (design 2 and design 3), the nanobody mutant VUN-XL (design 3) was designed. This final nanobody displayed the largest degree of potency and efficacy for a 5 residue CDR3-insertion of VUN701. The final VUN701 mutant was termed “VUN-XL” due to the addition of bulk into CDR3. In this nanobody, VUN701’s R103 is replaced by (GKAYSG).

### Orthosteric Volume Analysis

Orthosteric pocket volumes were quantified using POVME2^61^. For each GPCR of interest (ACKR3, MOR, B_2_AR, A_2A_R, and NT_1_R), all available experimental structures were retrieved from GPCRdb together with their corresponding conformational annotations (“active” or “inactive”). Experimental structures bound to an agonist, but carrying an “inactive” annotation were discarded. Individual PDB files were downloaded and processed using a custom python script that (i) removed all bound ligands, transducers, and non-protein atoms (ii) aligned each receptor to the ACKR3 reference structure, and (iii) combined each model into a single PDB file in which each model loaded as a single state of a multi-state file. After preprocessing, POVME2 was run on each structure with GridSpacing = 1 and DistanceCutoff = 1.6, with two inclusion spheres: one centered at (155.4, 156.7, 138.2), with a radius of 9, and the other centered at (155.1, 155.5, 133.8), with a radius of 8. The resulting volumes were plotted on a per-structure and state basis, and the complete dataset is provided in Supplementary Data 1.

### Data Visualization

All plots and structural visualizations were produced in GraphPad Prism, Schrodinger’s PyMOL, UCSF’s ChimeraX, or Python, with final figure assembly and curation in Adobe Illustrator.

## Acknowledgements

We thank Dr. Andrew Kleist, Dr. John Janetzko, Dr. John McCorvy, Dr. Raimond Heukers, and Dr. Christopher Schafer for their comments on the manuscript figures and text. We are grateful to Dr. Anthony Kossiakoff and Dr. Somnath Mukherjee for providing the NabFab and CID24 constructs used in this study.

This research was completed in part with computational resources and technical support provided by the Research Computing Center at the Medical College of Wisconsin.

This work was supported by a National Institutes of Health Allergy and Infectious Disease grant (NIAID R37AI058072 to BFV), by the Luxembourg Institute of Health (LIH) through the NanoLux Platform, the Cancer Foundation Luxembourg, Luxembourg National Research Fund (INTER/FWO Nanokine 15/10358798, PoC 19/14209621, INTER/FNRS CXCL12 20/15084569, CORE IMPACTT C23/BM/18068832, ACKROS C25/BM/19563724, INTER/AUDACE 25/19352183, PRIDE 21/16763386/CANBIO2), and by the European Union H2020-MSCA Program (Grant agreement 860229-ONCORNET2.0 for MJS).

## Author Contributions

RRS and BFV conceived the study and led all aspects of the project. RRS designed the experiments. RRS engineered and characterized nanobody variants. RRS, JD, AC, and MS performed functional signaling assays and conducted structure-guided mutagenesis. CBB, RRS, and ADLS prepared samples for cryo-EM, processed cryo-EM data, and determined and analyzed structures. LJO refined cryo-EM models. RRS, ADLS, SEJ, and FCP performed NMR experiments and analyzed spectroscopic data. RRS and SEJ carried out computational analyses, including volumetric measurements, comparative GPCR analyses, and literature analysis. BFV provided overall supervision, conceptual guidance, and critical input throughout the project. RRS, BFV, and AC contributed to experimental design, data interpretation, and manuscript editing. RRS and BFV wrote the manuscript. RRS created figures, with input from all authors. All authors discussed the results, edited the manuscript, and approved the final version of the manuscript.

