## Supplementary Materials for "Atypical GPCR Activation Resolved by Nanobody Engineering"

**This PDF file includes:**

Table S1: Endogenous GPCR interactions
Table S2: Characterization of all ACKR3 ligands
Table S3: Cryo-EM statistics table
Table S4: Characterization of nanobody-ACKR3 pharmacology
Table S5: Characterization of ACKR3 ECL2 mutant pharmacology
Table S6: Comparison of basal signaling in ACKR3 mutants
Table S7: Impact of "toggle-switch" mutations on CXCL12-induced ACKR3 activity
Table S8: Ligand-dependent effects of "toggle-switch" disruption on ACKR3 activity

Supplementary Figure 1: Pipeline for the structure determination and validation of the ACKR3–VUN701 complex
Supplementary Figure 2: Overview of VUN701 mutations to enable NabFab binding
Supplementary Figure 3: Comparison of ACKR3-VUN701 cryo-EM structure and model
Supplementary Figure 4: ACKR3-VUN701 density and unresolved regions
Supplementary Figure 5: Pipeline for the structure determination and validation of the ACKR3–VUN701 T105N complex.
Supplementary Figure 6: Full ACKR3 NMR spectra
Supplementary Figure 7: Pipeline for the structure determination and validation of the ACKR3–VUN-XL complex
Supplementary Figure 8: Validation of M138 as an NMR reporter of TM7 (Y315) movement in ACKR3
Supplementary Figure 9: Validation of I318M as a new NMR reporter of TM6 movement in ACKR3
Supplementary Figure 10: Representative views of orthosteric pocket volume measurements in ACKR3
Supplementary Figure 11: Ligand-dependent effects of "toggle-switch" disruption on ACKR3 activity

Supplementary Movie 1: 3DVA of ACKR3-VUN701

Supplementary Data 1: Summary of GPCR orthosteric pocket volume calculations

| GPCR | Ligand | Family | Size (Da) | Source (PMID) |
|---|---|---|---|---|
| ACKR3 | DynorphinA | Opioid | 2147.51 | 32561830 |
| | DynorphinA 1-13 | Opioid | 1604.00 | 32561830 |
| | DynorphinB | Opioid | 1570.86 | 32561830 |
| | Leumorphin | Opioid | 3351.64 | 32561830 |
| | Big Dynorphin | Opioid | 3984.72 | 32561830 |
| | Adrenorphin | Opioid | 985.17 | 32561830 |
| | BAM22 | Opioid | 2839.25 | 32561830 |
| | fΨg nociceptin 1-13 | Opioid | 1178.36 | 32561830 |
| | Adrenomedullin | Proadrenomedullin-dervied peptide | 6031.80 | 33860204 |
| | PAMP-12 | Proadrenomedullin-dervied peptide | 1619.94 | 33860204 |
| | CXCL11 | Chemokine | 8303.00 | 16940167 |
| | CXCL12 | Chemokine | 7949.40 | 16107333 |
| | vCCL2 | Chemokine | 8397.88 | 27238288 |
| | MIF | Cytokine | 12345.11 | 25266363 |
| μOR | b-endorphin | Opioid | 3465.02 | 999687 |
| | Met-enkephalin | Opioid | 573.66 | 1207728 |
| | Leu-enkephalin | Opioid | 555.63 | 1207728 |
| | Dynorphin a | Opioid | 2147.51 | 230519 |
| $\beta_2$AR | Adrenaline | Catecholamines | 183.207 | 18882199 |
| | Noradrenaline | Catecholamines | 169.18 | 18882199 |
| $A_{2A}$R | Adenosine | Nucleotide | 267.245 | 2125216 |
| $NT_1$R | Neurotensin | Neuropeptide | 1672.92 | 1167549 |
| | Neuromedin N | Neuropeptide | 745.949 | 3019713 |

**Table S1: Endogenous GPCR interactions.**

Endogenous ligands for ACKR3 are shown alongside ligands for μ-opioid receptor (μOR), $\beta_2$-adrenergic receptor ($\beta_2$AR), adenosine $A_{2A}$ receptor ($A_{2A}$R), and neurotensin receptor 1 ($NT_1$R) for comparison. For each ligand, the signaling family, the molecular size (Da), and the primary literature source in which the interaction was first described (PMID) are indicated.

| Ligand | Ligand Type | Endogenous | Pharmacology | Identifying Source (PMID) |
|---|---|---|---|---|
| CXCL11 | Chemokine | Yes | agonist | 16940167 |
| CXCL12 | Chemokine | Yes | agonist | 16107333 |
| vCCL2 | Chemokine | Yes | agonist | 27238288 |
| Dynorphin A | Opioid Peptide | Yes | agonist | 32561830 |
| Dynorphin A 1-13 | Opioid Peptide | Yes | agonist | 32561830 |
| Dynorphin B | Opioid Peptide | Yes | agonist | 32561830 |
| Leumorphin | Opioid Peptide | Yes | agonist | 32561830 |
| Big Dynorphin | Opioid Peptide | Yes | agonist | 32561830 |
| Adrenorphin | Opioid Peptide | Yes | agonist | 32561830 |
| BAM22 | Opioid Peptide | Yes | agonist | 32561830 |
| FΨG nociceptin 1-13 | Opioid Peptide | Yes | agonist | 32561830 |
| ADM | Peptide Hormone | Yes | agonist | 33860204 |
| PAMP-12 | Peptide Hormone | Yes | agonist | 33860204 |
| MIF | Cytokine | Yes | agonist | 25266363 |
| Conolidine | Small Molecule | No | agonist | 34075018 |
| CCX777 | Small Molecule | No | agonist | 28098154 |
| CCX771 | Small Molecule | No | agonist | 19641136 |
| CCX754 | Small Molecule | No | agonist | 19001056 |
| CCX733 | Small Molecule | No | agonist | 19001056 |
| CCX662 | Small Molecule | No | agonist | 31506383 |
| TC14012 | Small Molecule | No | agonist | 20956518 |
| AMD3100 | Small Molecule | No | agonist | 19255243 |
| VUF15485 | Small Molecule | No | agonist | 38346795 |
| VUF25444 | Small Molecule | No | agonist | 38732237 |
| VUF16545 | Small Molecule | No | agonist | 23333329 |
| VUF11403 | Small Molecule | No | agonist | 22424612 |
| VUF11207 | Small Molecule | No | agonist | 22424612 |
| NUCC-54129 | Small Molecule | No | agonist | 35809897 |
| NUCC-200823 | Small Molecule | No | agonist | 35809897 |
| PF-06827080 [Compound 18] | Small Molecule | No | agonist | 29627981 |
| LIH383 | Peptide | No | agonist | 32561830 |
| Nb1 | Nanobody | No | antagonist | 23979133 |
| Nb2 | Nanobody | No | antagonist | 23979133 |
| Nb3 | Nanobody | No | antagonist | 23979133 |
| Compound 10 | Small Molecule | No | antagonist | 32551020 |
| VUN701 | Nanobody | No | antagonist | 35857540 |
| ACT-1004-1239 | Small Molecule | No | antagonist | 33314938 |
| VUF16480 | Small Molecule | No | inverse agonist | 41317408 |
| VUN700 | Nanobody | No | inverse agonist | 39574661 |
| VUN702 | Nanobody | No | inverse agonist | 39574661 |
| LN6023 | Small Molecule | No | superagonist | 36150079 |
| LN5972 | Small Molecule | No | superagonist | 36150079 |

**Table S2: Characterization of all ACKR3 ligands.**

All endogenous and synthetic molecules reported to modulate ACKR3 activity. Molecules are listed according to ligand type, described pharmacology, and the primary literature source in which the interaction was first described (PMID).

| **Maps** | | | | | |
|---|---|---|---|---|---|
| Description | Global map | Local map | Local map | Global map | Global map |
| Sample | **ACKR3-VUN701 NabFab AntiFabNb** | **ACKR3-VUN701 NabFab** | **ACKR3-VUN701** | **ACKR3-VUN-XL NabFab AntiFabNb CID24 AntiFabNb** | **ACKR3-VUN701 T105N NabFab AntiFabNb** |
| EMDB: | (EMDB-XXXXX) | | | (EMDB-XXXXX) | (EMDB-XXXXX) |
| RCSB PDB: | (PDB XXXX) | | | (PDB XXXX) | (PDB XXXX) |
| **Collection** | | | | | |
| Magnification | 105,000 | 105,000 | 105,000 | 165,000 | 165,000 |
| Voltage (kV) | 300 | 300 | 300 | 200 | 200 |
| Electron exposure (e-$^{2}$/Å) | 50.3 | 50.3 | 50.3 | 60.0 | 60.0 |
| Defocus range (μm) | -0.8 to -2.0 | -0.8 to -2.0 | -0.8 to -2.0 | -1.0 to -3.0 | -1.0 to -3.0 |
| Pixel size (Å) | 0.85 (physical) | 0.85 (physical) | 0.85 (physical) | 0.71 (physical) | 0.71 (physical) |
| Symmetry imposed | C1 | C1 | C1 | C1 | C1 |
| Initial particle images (no.) | 12,271,852 | 12,271,852 | 12,271,852 | 3,186,525 | 10,989,551 |
| Final particle images (no.) | 39,498 | 39,498 | 39,498 | 146,437 | 361,678 |
| Map resolution (Å) (masked) | 3.3 | 3.2 | 3.5 | 3.9 | 3.4 |
| FSC threshold | 0.143 | 0.143 | 0.143 | 0.143 | 0.143 |
| **Refinement** | | | | | |
| Map sharpening B factor (Å$^{2}$) | 71.9 | 75.3 | 85.7 | 144.4 | 162.9 |

**Table S3: Cryo-EM statistics table.**

Summary of statistics for cryo-EM maps generated in this study.

| Ligand | log($EC_{50}$) | Span |
|---|---|---|
| CXCL12 | -9.64 ± 0.06 | 0.140 ± 0.004 |
| VUN701 | N/A | N/A |
| VUN701 T105A | -6.78 ± 0.35 | 0.015 ± 0.003 |
| VUN701 T105N | -7.11 ± 0.32 | 0.023 ± 0.003 |
| VUN701 T105W | -8.05 ± 0.35 | 0.019 ± 0.003 |
| VUN701 T105Q | -8.13 ± 0.20 | 0.028 ± 0.002 |
| VUN701 T105K | -7.05 ± 0.17 | 0.036 ± 0.003 |
| VUN-XL | -6.25 ± 0.08 | 0.098 ± 0.003 |
| VUN-XL #2 | -6.46 ± 0.10 | 0.069 ± 0.003 |
| VUN-XL #3 | -5.32 ± 0.10 | 0.074 ± 0.004 |

**Table S4: Characterization of nanobody-ACKR3 pharmacology.**
Summary of $EC_{50}$ and Span (maximum signaling amplitude) values for CXCL12, VUN701, or VUN701 mutants in a BRET-based β-arrestin2 recruitment assay with WT ACKR3. N = 3 biologically independent experiments plotted as mean +/- SEM.

| Receptor | log($EC_{50}$) | Span | Basal Level |
|---|---|---|---|
| WT ACKR3 | -9.73 ± 0.09 | 0.99 ± 0.04 | 0.01 ± 0.03 |
| ACKR3 $R197E^{45x51}$ | N/A | 0.09 ± 0.05 | -0.29 ± 0.05 |
| ACKR3 $E202R^{ECL2}$ | N/A | 0.08 ± 0.04 | 0.93 ± 0.03 |

**Table S5: Characterization of ACKR3 ECL2 mutant pharmacology.**
Summary of $EC_{50}$, Span (maximum signaling amplitude), and basal signaling levels (Fold Change vs WT) for CXCL12 in a BRET-based β-arrestin2 recruitment assay with WT ACKR3, ACKR3 $R197E^{45x51}$, and ACKR3 $E202R^{ECL2}$. N = 3 biologically independent experiments plotted as mean +/- SEM.

| ACKR3 Construct | Basal BRET Response |
|---|---|
| irrDNA | 0.223 ± 0.003 |
| ST/A | 0.252 ± 0.012 |
| WT | 0.313 ± 0.029 |
| R142A$^{3x50}$ | 0.266 ± 0.013 |
| R142E $^{3x50}$ | 0.261 ± 0.015 |
| Y232A$^{5x58}$ | 0.245 ± 0.002 |
| Y232L$^{5x58}$ | 0.360 ± 0.010 |
| D244A$^{ICL3}$ | 0.939 ± 0.071 |
| D244K$^{ICL3}$ | 0.827 ± 0.067 |
| E246A$^{6x29}$ | 0.772 ± 0.044 |
| E246K$^{6x29}$ | 0.955 ± 0.035 |
| K247A$^{6x30}$ | 0.264 ± 0.017 |
| K247E $^{6x30}$ | 0.257 ± 0.012 |
| Y257A$^{6x40}$ | 0.291 ± 0.017 |
| Y257L$^{6x40}$ | 0.271 ± 0.021 |
| Y315A$^{7x53}$ | 0.242 ± 0.001 |
| Y315L$^{7x53}$ | 0.253 ± 0.022 |

**Table S6: Comparison of basal signaling in ACKR3 mutants.**
Summary of basal signaling levels for ACKR3 in a nanoBRET-based β-arrestin2 recruitment assay with no added ligand. N = 3 biologically independent experiments plotted as mean +/- SEM.

| ACKR3 Construct | log(EC50) (M) | Span |
|---|---|---|
| WT | -9.12 ± 0.10 | 0.268 ± 0.013 |
| W265Q$^{6x48}$ | -8.83 ± 0.12 | 0.184 ± 0.010 |
| W265F$^{6x48}$ | -8.48 ± 0.21 | 0.195 ± 0.020 |
| W265A$^{6x48}$ | -8.82 ± 0.11 | 0.213 ± 0.011 |

**Table S7: Impact of "toggle-switch" mutations on CXCL12-induced ACKR3 activity.** Summary of $EC_{50}$ and Span (maximum signaling amplitude) for CXCL12 in a nanoBRET-based β-arrestin2 recruitment assay with WT ACKR3 and the ACKR3 "toggle-switch" mutants W265Q$^{6x48}$, W265F$^{6x48}$, and W265A$^{6x48}$. N = 3 biologically independent experiments plotted as mean +/- SEM.

| Ligand | WT | | W265A$^{6x48}$ | | Toggle Switch Required |
|---|---|---|---|---|---|
| | log(EC50) (M) | Span | log(EC50) (M) | Span | |
| CXCL12 | -8.58 ± 0.11 | 0.061 ± 0.003 | -8.19 ± 0.10 | 0.084 ± 0.004 | No |
| Chimera$_{X12\text{-}X11}$ | -7.97 ± 0.15 | 0.053 ± 0.004 | -7.51 ± 0.24 | 0.074 ± 0.009 | No |
| VUF25444 | -8.65 ± 0.08 | 0.093 ± 0.004 | -6.73 ± 0.20 | 0.089 ± 0.010 | No |
| VUF16545 | -8.83 ± 0.24 | 0.076 ± 0.012 | -7.31 ± 0.34 | 0.056 ± 0.009 | No |
| LIH366 | -7.22 ± 0.169 | 0.076 ± 0.006 | -5.41 ± 0.13 | 0.051 ± 0.004 | No |
| CXCL11 | -8.16 ± 0.11 | 0.048 ± 0.003 | -7.28 ± 0.33 | 0.014 ± 0.003 | Yes |
| vCCL2 | -7.83 ± 0.11 | 0.049 ± 0.003 | N/A | N/A | Yes |
| BAM22 | -6.97 ± 0.07 | 0.063 ± 0.002 | N/A | N/A | Yes |
| PAMP12 | -5.85 ± 0.15 | 0.074 ± 0.009 | N/A | N/A | Yes |
| Dynorphin A | -6.75 ± 0.12 | 0.071 ± 0.005 | N/A | N/A | Yes |
| LIH383 | -7.56 ± 0.06 | 0.060 ± 0.002 | N/A | N/A | Yes |
| VUF11207 | -7.12 ± 0.16 | 0.082 ± 0.006 | N/A | N/A | Yes |
| VUF15485 | -7.50 ± 0.11 | 0.094 ± 0.005 | N/A | N/A | Yes |

**Table S8: Ligand-dependent effects of “toggle-switch” disruption on ACKR3 activity.** Summary of $EC_{50}$ and Span (maximum signaling amplitude) for a large panel of ACKR3 ligands in a NanoBRET-based β-arrestin2 recruitment assay with WT ACKR3 or the ACKR3 “toggle-switch” mutant W265A$^{6x48}$. N/A indicates no measurable β-arrestin2 recruitment. N = 3 biologically independent experiments plotted as mean +/- SEM.

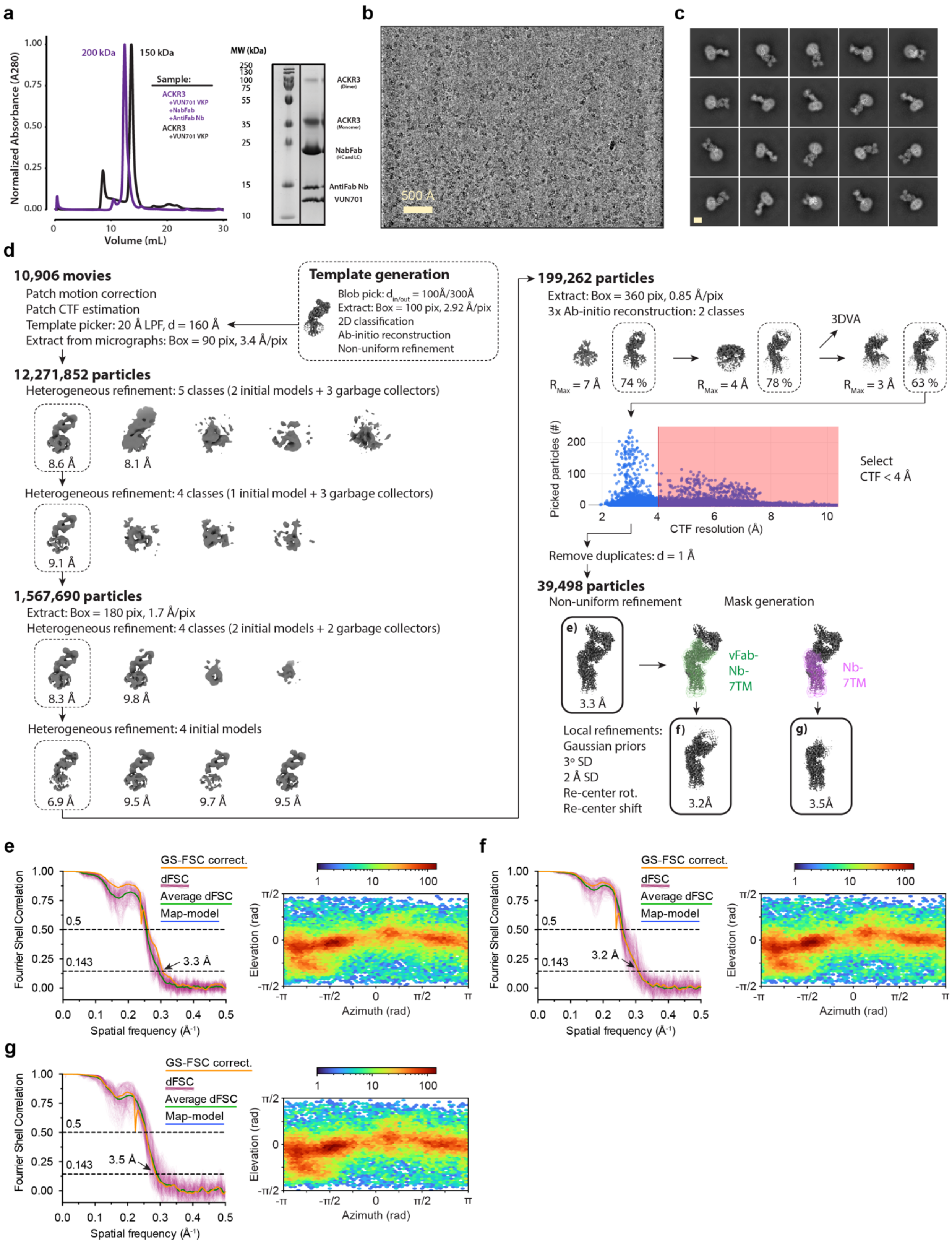


**Supplementary Figure 1: Pipeline for the structure determination and validation of the ACKR3–VUN701 complex.**

a) Left: Size-exclusion chromatography profile of the purified ACKR3-VUN701 complex alone (black), and with the addition of the NabFab and AntiFabNb (purple). Right: SDS–PAGE showing ACKR3-VUN701-NabFab-AntiFabNb sample composition and purity. Representative (b) cryo-EM micrograph of vitrified particles collected for structure determination and (c) representative 2D-class averages are shown to the right. d) Cryo-EM image processing workflow. An initial dataset of 10,906 movies was motion-corrected and CTF-estimated, followed by particle picking, and successive rounds of heterogeneous refinement to remove poorly aligned particles and junk classes. Final non-uniform refinement yielded a high-quality reconstruction from 39,498 particles. (e–g) Fourier shell correlation (FSC) curves and directional FSC analyses for the final reconstructions deposited in the EMDB. The final map resolutions were determined using the 0.143 FSC criterion (indicated).

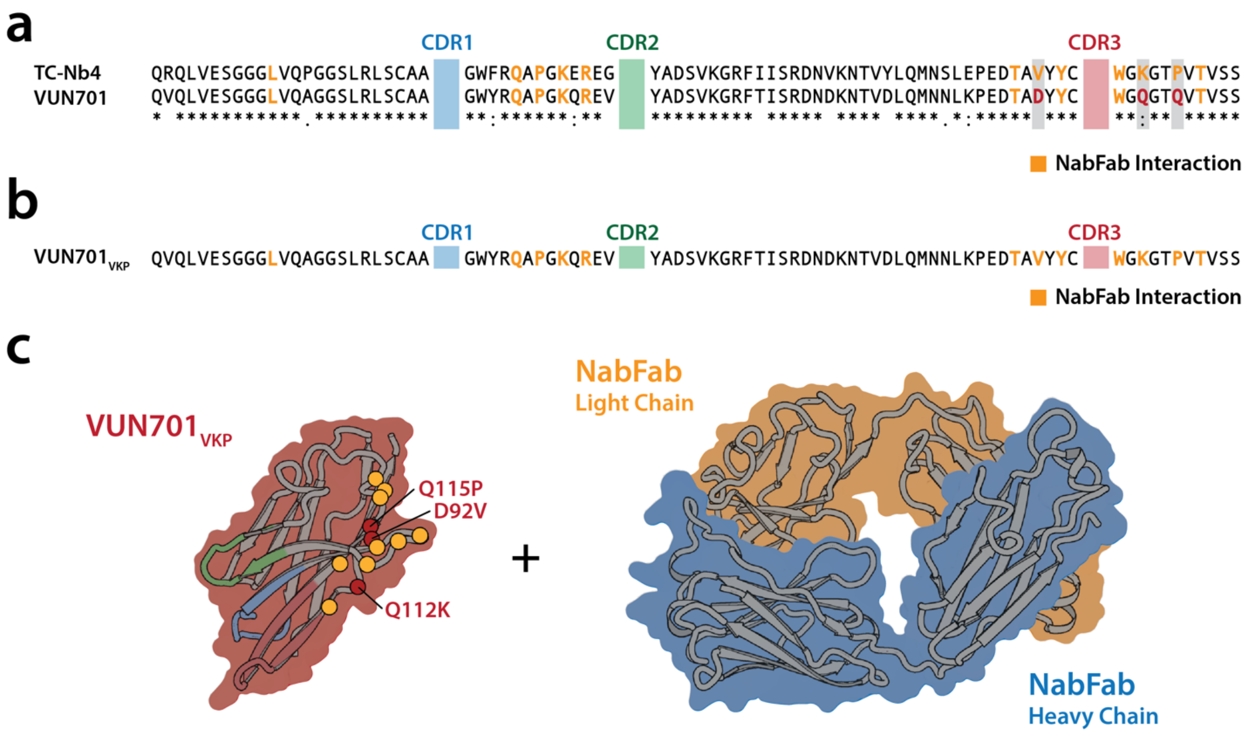


**Supplementary Figure 2: Overview of VUN701 mutations to enable NabFab binding.** a) Sequence alignment of the alpaca nanobody TC-Nb4, which was used to generate the nanobody-binding Fab (NabFab), with the llama-generated VUN701. All CDR regions are removed in this alignment. Residues that facilitate NabFab binding are marked as gold, bolded text. VUN701 differs in these conserved regions at D92, Q112, and Q115 (red). b) Sequence of $VUN701_{VKP}$ mutant, which contains the mutations D92V, Q112K, and Q115P to facilitate NabFab binding. c) Cartoon of $VUN701_{VKP}$ and the NabFab illustrating that NabFab binding occurs away from the CDR interactions.

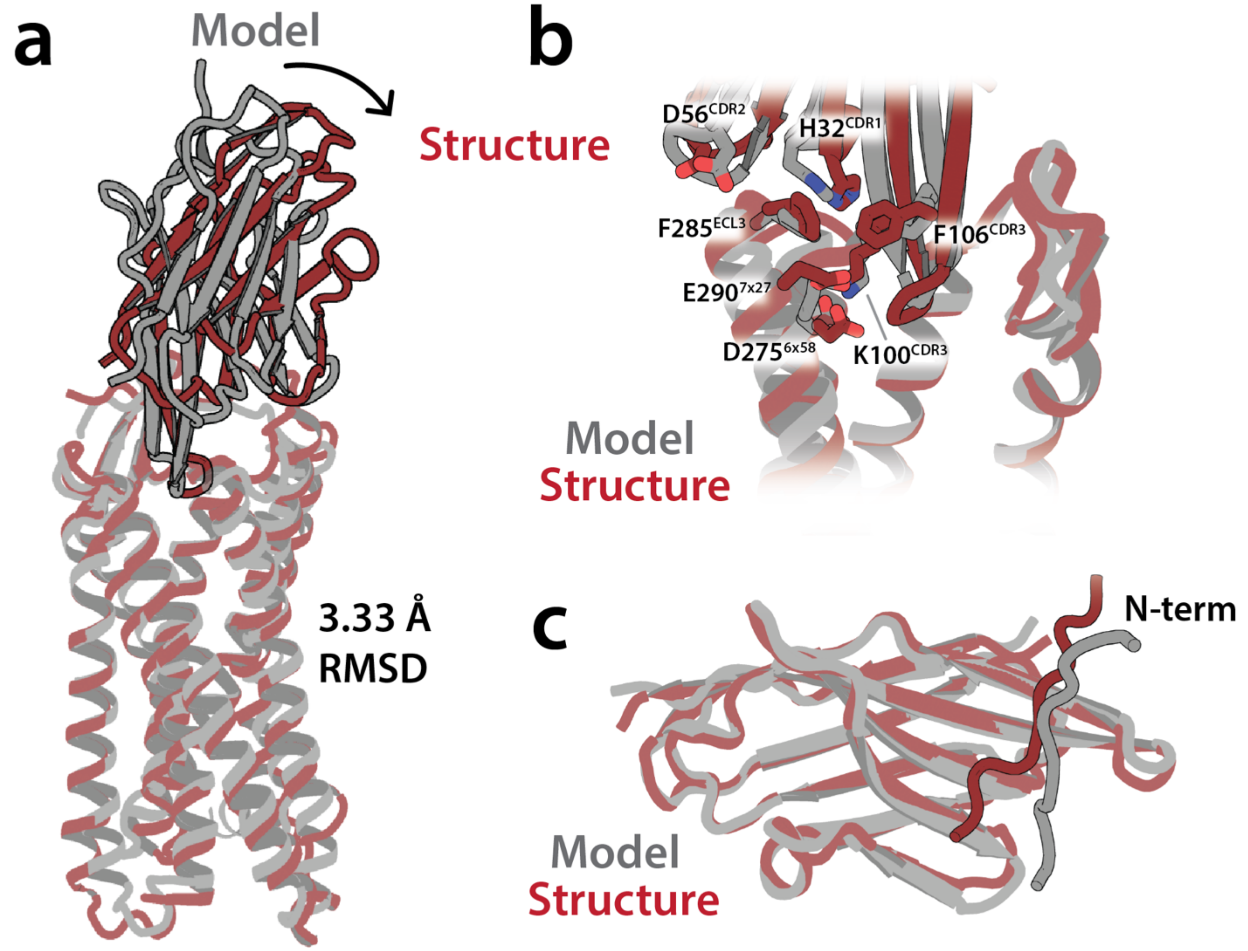


**Supplementary Figure 3: Comparison of ACKR3-VUN701 cryo-EM structure and model.** Overlay of the ACKR3-VUN701 cryo-EM structure with the previously described ACKR3-VUN701 model. a) A 3.33 Angstrom total backbone RMSD was calculated between the two structures. Differences in structure primarily occur in the VUN701 binding pose which was more upright in the ACKR3-VUN701 model. b) All previously described contacts between ACKR3 and VUN701 in the extracellular space were identical. c) Contacts between VUN701 and ACKR3's N-terminus were additionally confirmed, though these were not tested previously.

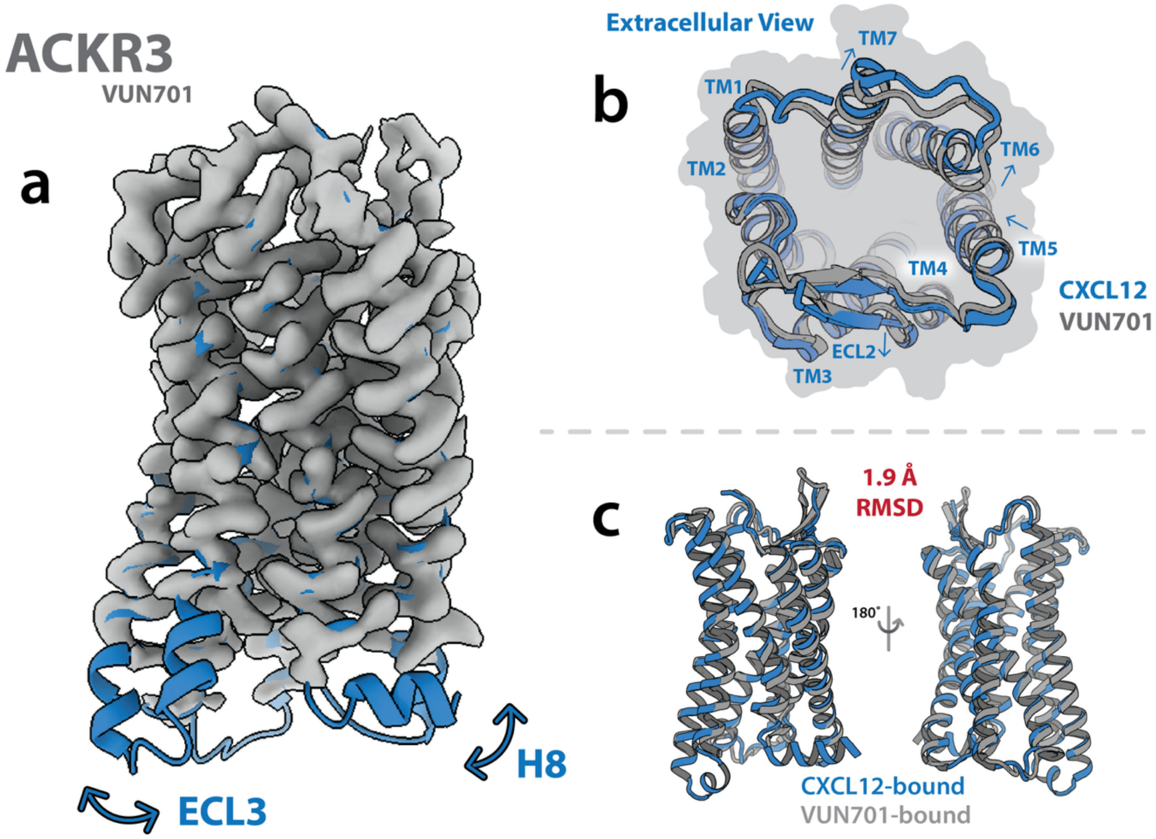


**Supplementary Figure 4: ACKR3-VUN701 density and unresolved regions.**
a) Cryo-EM density map of ACKR3 bound to the neutral antagonist nanobody VUN701, shown as a surface representation over a modified structural model (PDB ID: XXXX). Intracellular regions (TM5, TM6, TM7, and helix 8) are shown for visualization but are poorly resolved or absent from the map, reflecting substantial conformational heterogeneity in these regions. b) Extracellular view of ACKR3 comparing the VUN701-bound structure (gray) with the CXCL12-bound active-state structure (blue; PDB: 7SK5). The extracellular helical arrangement and ECL2 conformation are highly similar between the two states. c) Structural overlay of VUN701-bound ACKR3 (gray) with the active-state ACKR3-CXCL12 complex (blue) reveals that VUN701 stabilizes a pseudo-active conformation despite antagonizing receptor activity.

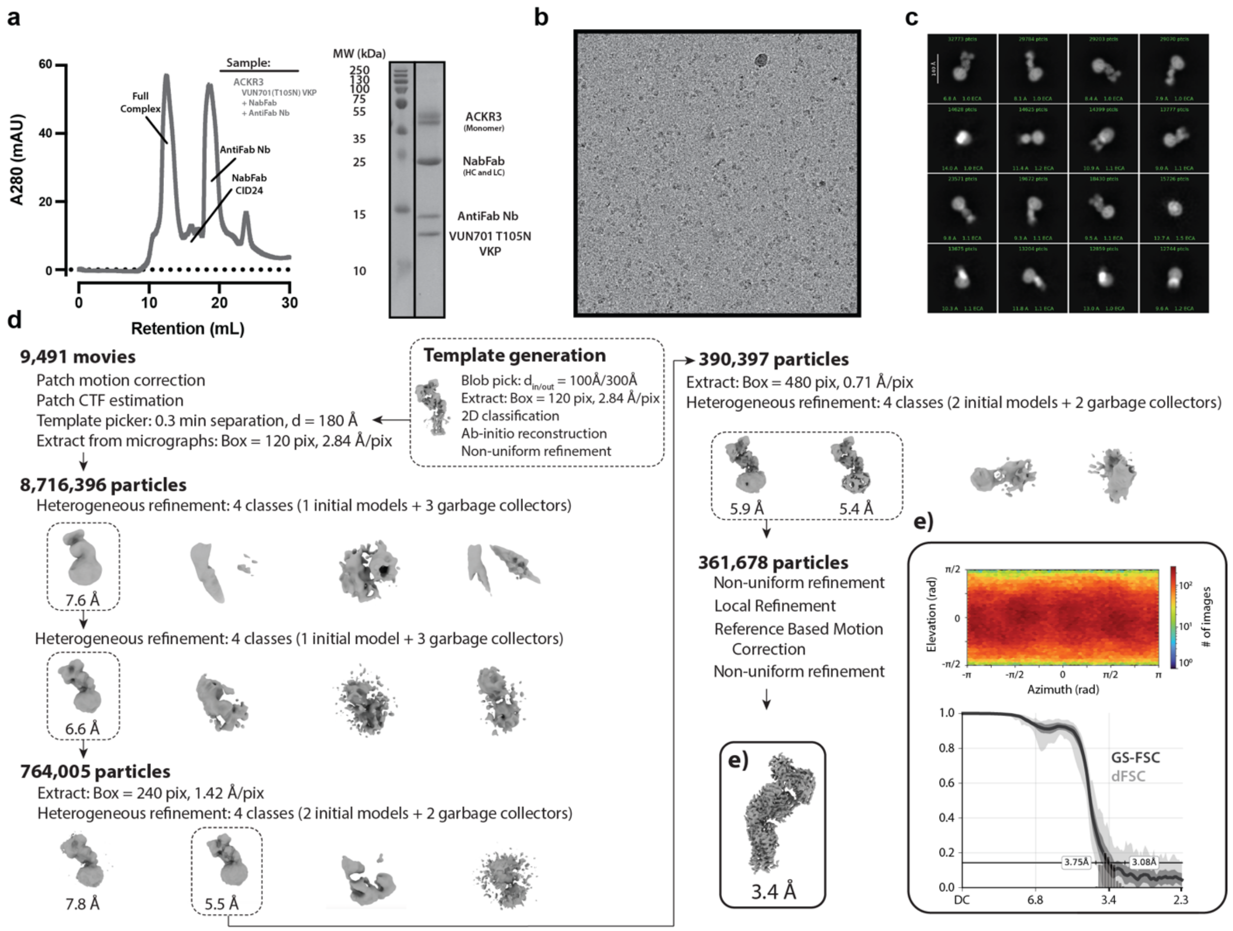


**Supplementary Figure 5: Pipeline for the structure determination and validation of the ACKR3–VUN701 T105N complex.**

a) Left: Size-exclusion chromatography profile of the complexed ACKR3-VUN701(T105N)-NabFab-AntiFabNb. Right:  SDS–PAGE showing ACKR3-VUN701(T105N)-NabFab-AntiFabNb sample composition and purity. Representative (b) cryo-EM micrograph of vitrified particles collected for structure determination, and (c) representative 2D-class averages are shown to the right. d) Cryo-EM image processing workflow. An initial dataset of 9,491 movies was motion-corrected and CTF-estimated, followed by particle picking, and successive rounds of heterogeneous refinement to remove poorly aligned particles and junk classes. Final non-uniform refinement yielded a high-quality reconstruction from 361,678 particles. (e) The Fourier shell correlation (FSC) curve and directional FSC analysis for the final reconstructions deposited in the EMDB. The final map resolutions were determined using the 0.143 FSC criterion (indicated).

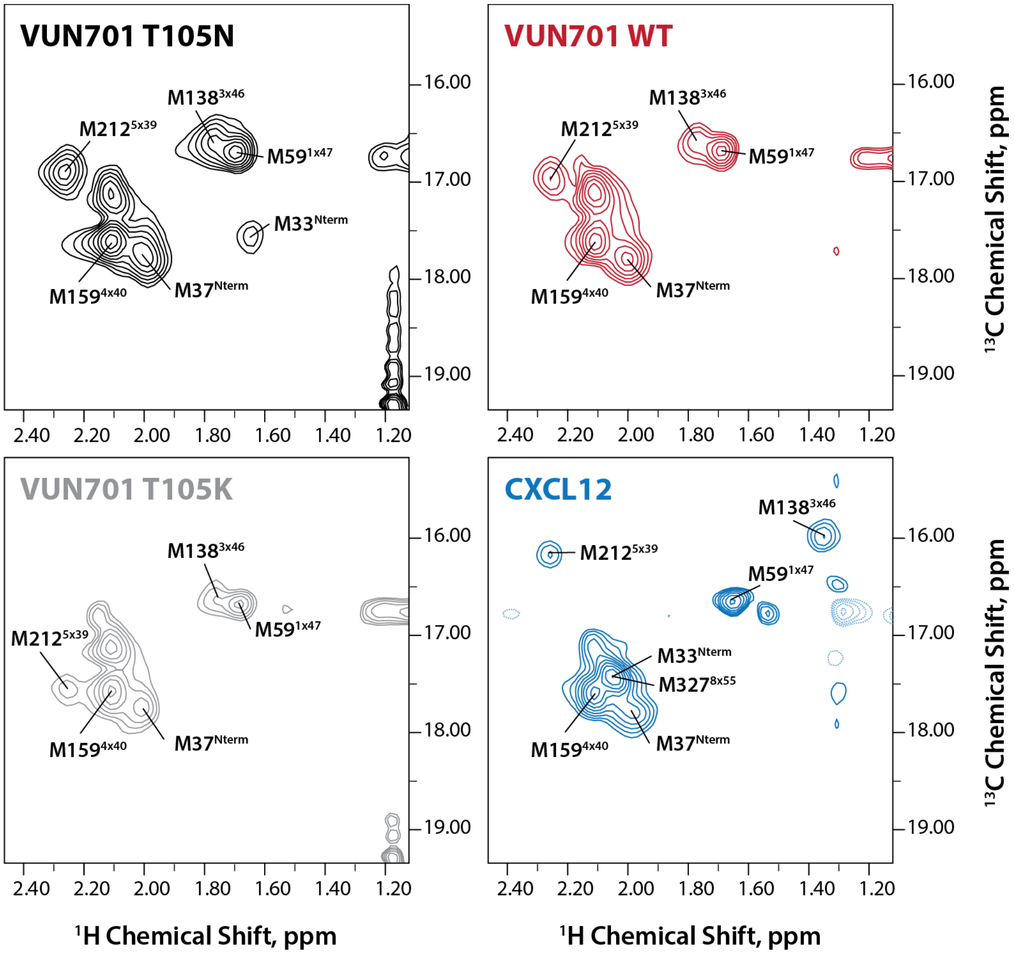


**Supplementary Figure 6: Full ACKR3 NMR spectra.**
$^{1}$H-$^{13}$C heteronuclear single quantum coherence NMR spectra of $^{13}$C-ε-Methionine labeled WT-ACKR3 with various ligands at 310 K. Assigned Met residues are labeled.

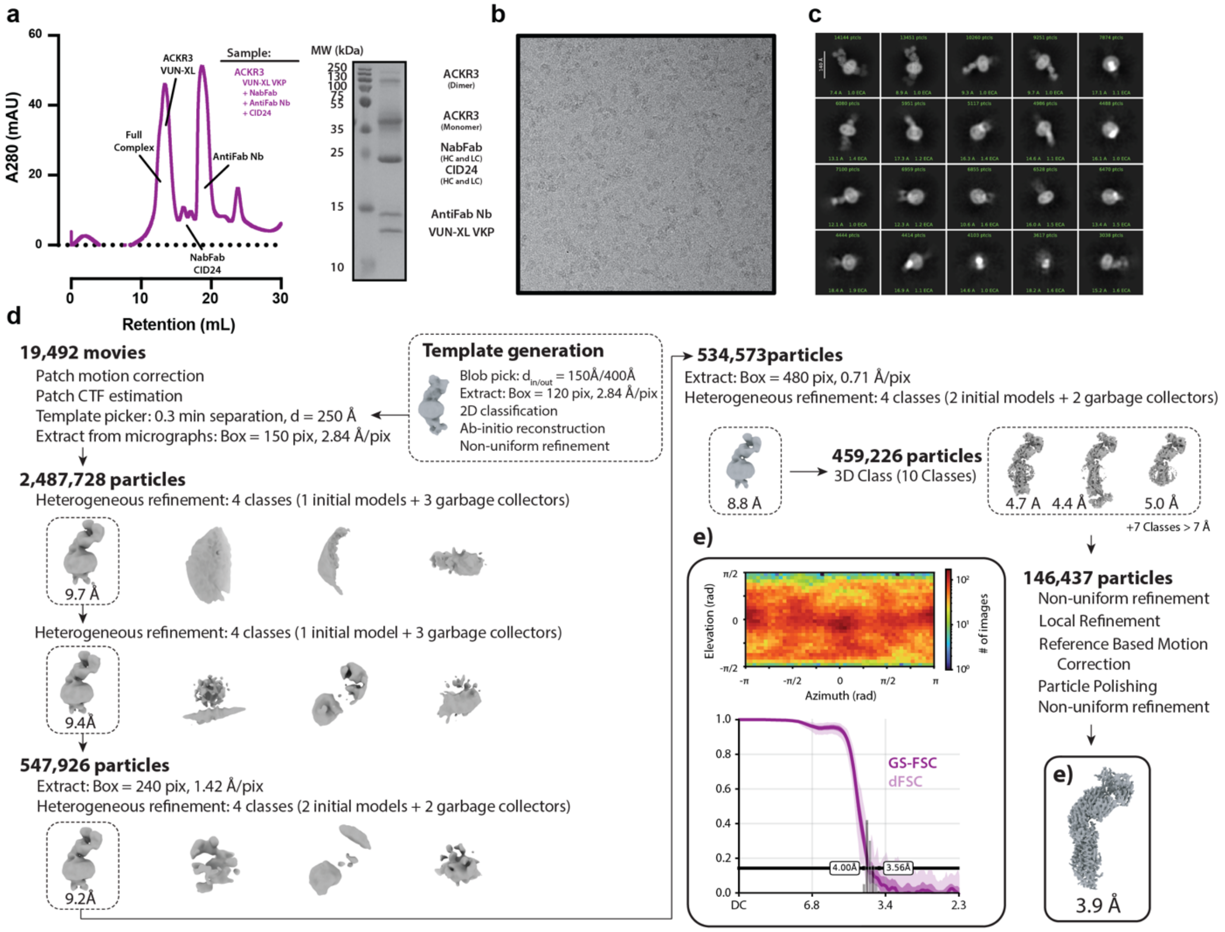


**Supplementary Figure 7: Pipeline for the structure determination and validation of the ACKR3–VUN-XL complex.**

a) Left: Size-exclusion chromatography profile of the complexed ACKR3-VUNXL-NabFab-CID24-AntiFabNb. Right: SDS–PAGE showing ACKR3-VUN701XL-NabFab-CID24-AntiFabNb sample composition and purity. Representative (b) cryo-EM micrograph of vitrified particles collected for structure determination, and (c) representative 2D-class averages are shown to the right. d) Cryo-EM image processing workflow. An initial dataset of 19,492 movies was motion-corrected and CTF-estimated, followed by particle picking, and successive rounds of heterogeneous refinement to remove poorly aligned particles and junk classes. 3D-class averages, followed by final non-uniform refinement yielded a high-quality reconstruction from 146,437 particles. (e) The Fourier shell correlation (FSC) curve and directional FSC analysis for the final reconstructions deposited in the EMDB. The final map resolutions were determined using the 0.143 FSC criterion (indicated).

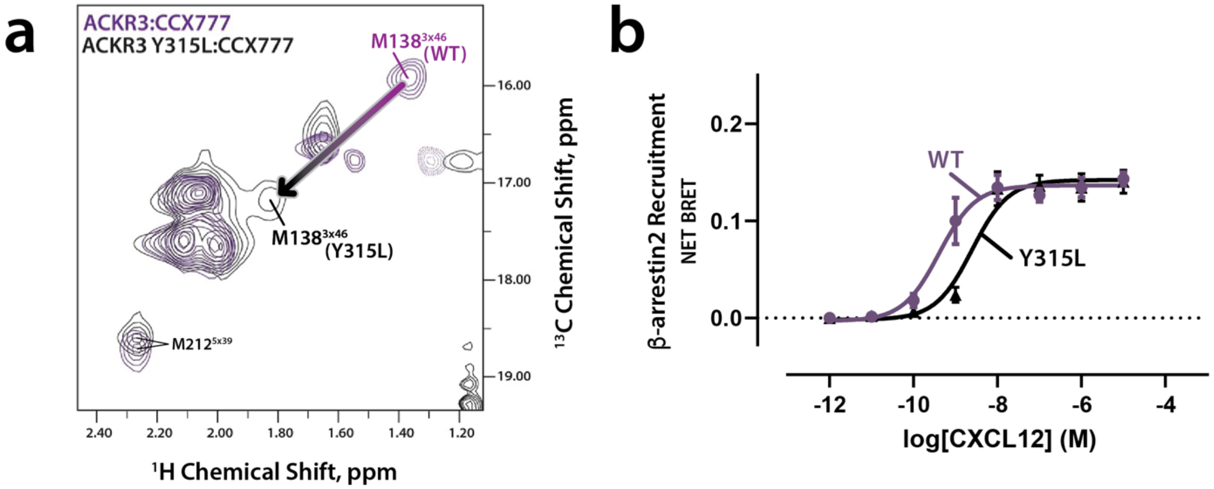


**Supplementary Figure 8: Validation of M138 as an NMR reporter of TM7 (Y315) movement in ACKR3.**

a) $^{1}H$-$^{13}C$ heteronuclear single quantum coherence NMR spectra of $^{13}C$-ε-Methionine labeled WT-ACKR3 overlayed with the spectra of Y315L ACKR3. In each spectrum, ACKR3 is bound to the small molecule agonist CCX777 as previously described[18]. In the wild-type receptor, M138 exhibits a pronounced chemical shift consistent with an aromatic ring-current effect arising from proximity to $Y315^{7x53}$. Substitution of Y315 with leucine (Y315L) abolishes this shift, demonstrating that the M138 signal directly reports on the position of Y315 and, by extension, TM7 movement. b) BRET-based β-arrestin2 recruitment assay of WT ACKR3 or ACKR3 Y315L with endogenous ligand CXCL12 demonstrating that despite the loss of the M138 NMR signature, the Y315L mutation preserves agonist-induced signaling and conformational changes. N = 3 biologically independent experiments plotted as mean +/- SEM.

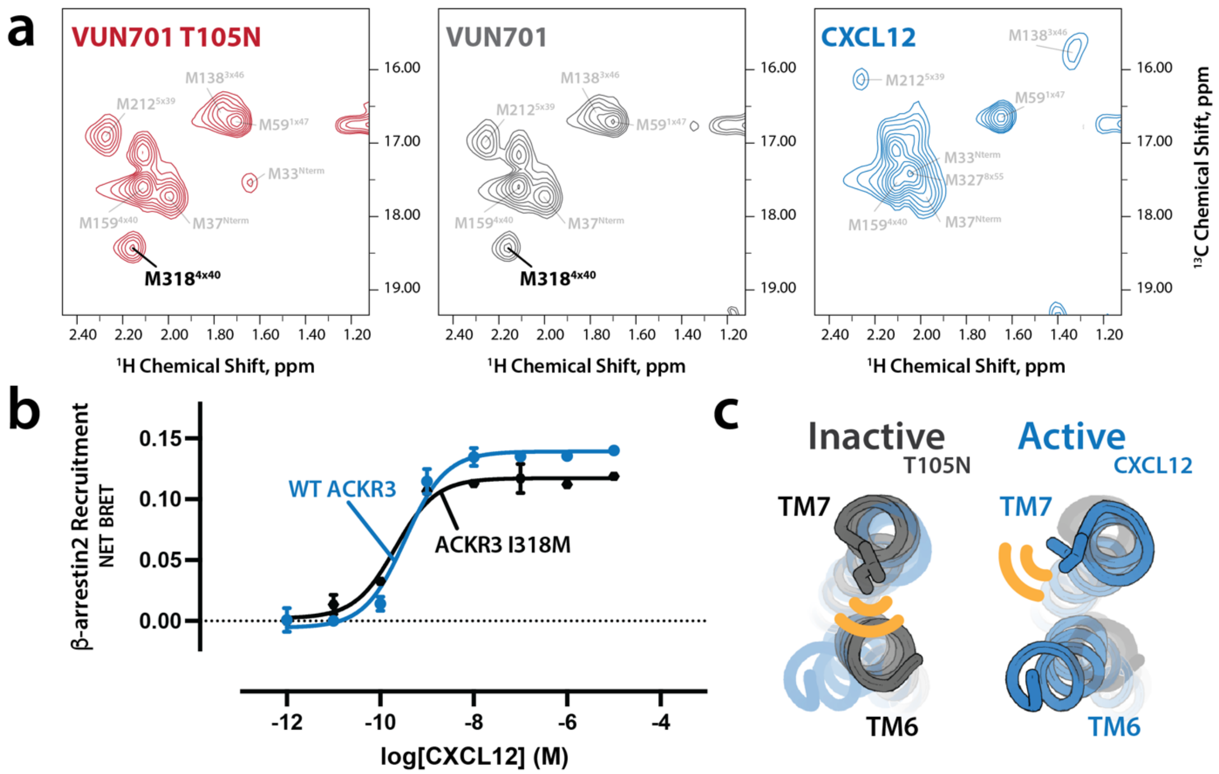


**Supplementary Figure 9: Validation of I318M as a new NMR reporter of TM6 movement in ACKR3.**

a) $^{1}H$-$^{13}C$ heteronuclear single quantum coherence (HSQC) NMR spectra of ACKR3 I318M in complex with CXCL12 (agonist), VUN701 (neutral antagonist), or VUN701 T105N (inverse agonist). In the inactive receptor state (VUN701, VUN701 T105N), I318M packs against TM6, constraining the side chain in a defined rotameric state and giving rise to a sharp resonance at ~18.3 ppm ($^{13}C$). In the CXCL12-bound active conformation, outward displacement of TM6 relieves this packing interaction, rendering I318M conformationally unrestricted and causing the resonance to merge with otherMet peaks with random-coil chemical shifts (~17 ppm $^{13}C$, 2.1 ppm $^{1}H$). b) β-arrestin2 recruitment measured by BRET for WT ACKR3 and the I318M mutant in response to CXCL12. The I318M substitution preserves agonist-induced arrestin recruitment, demonstrating that introduction of the NMR probe does not perturb receptor activation while providing a sensitive readout of TM6 conformational dynamics. c) Structural illustration of the active and inactive states of ACKR3, demonstrating the change in chemical environment around I318 in response to activation. N = 3 biologically independent experiments plotted as mean +/- SEM.

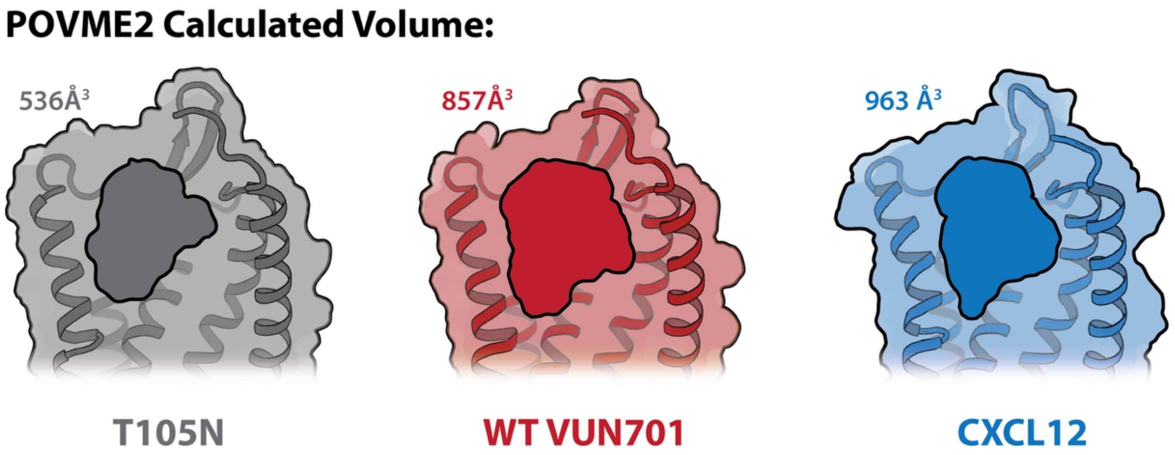


**Supplementary Figure 10: Representative views of orthosteric pocket volume measurements in ACKR3.**

Illustration of the orthosteric volume calculated for ACKR3 using POVME2 in the inactive state (VUN701 T105N), intermediate state (VUN701), and active state (CXCL12).

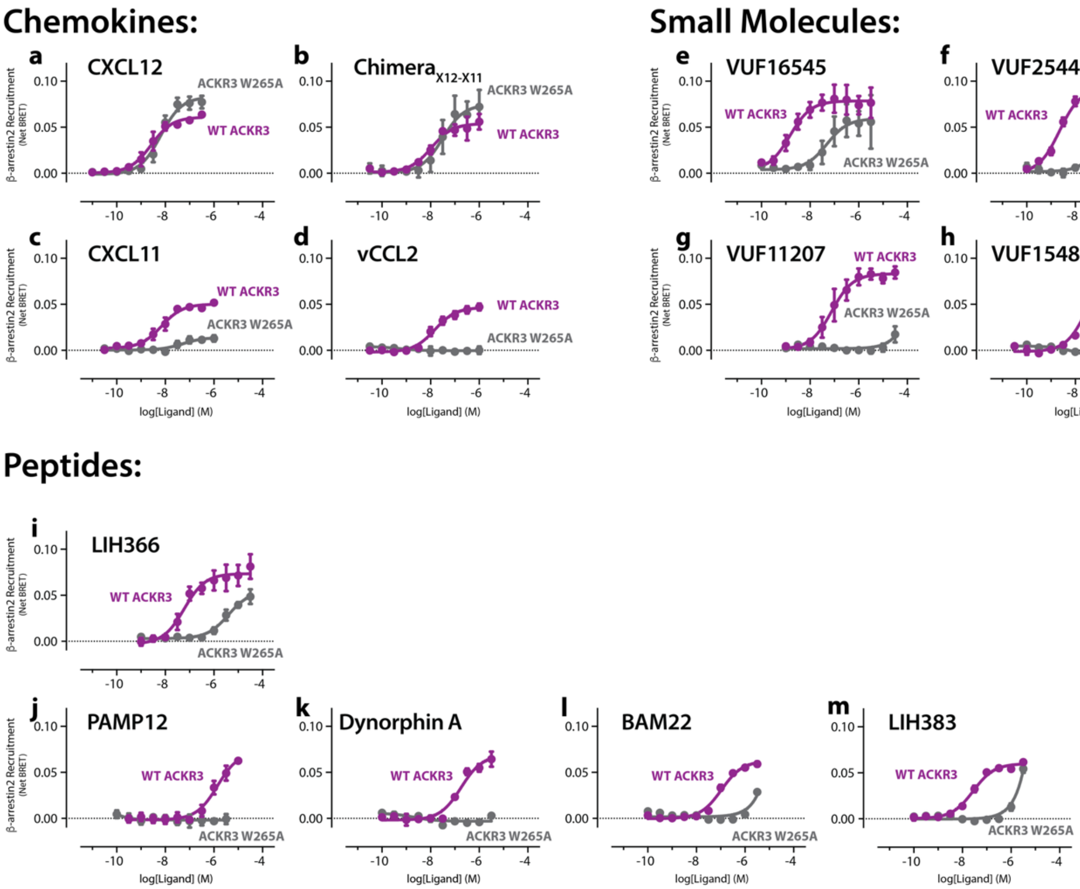


**Supplementary Figure 11: Ligand-dependent effects of "toggle-switch" disruption on ACKR3 activity.**

NanoBRET-based β-arrestin2 recruitment assay with WT ACKR3 (purple) or the ACKR3 "toggle-switch" mutant W265A$^{6x48}$ (gray) for the ACKR3 ligands CXCL12 (a), Chimera$_{X12\text{-}X11}$ (b), CXCL11 (c), vCCL2 (d), VUF16545 (e), VUF25444 (f), VUF11207 (g), VUF15485 (h), LIH366 (i), PAMP12 (j), dynorphin A (k), BAM22 (l), and LIH383 (m). N = 3 biologically independent experiments plotted as mean +/- SEM.

**Supplementary Movie 1: 3DVA of ACKR3-VUN701**

Three-dimensional variability analysis (3DVA) of the ACKR3–VUN701 cryo-EM complex, illustrating coordinated conformational changes in ECL2, TM5, TM6, and TM7 captured along the primary mode of variability (principal component 0).

**Supplementary Data 1: Summary of GPCR orthosteric pocket volume calculations.** The dataset includes, for each structure analyzed, the receptor name, PDB ID, activation state, bound ligand, and calculated orthosteric pocket volume ($Å^3$).
